# Regulatory divergence and effector turnover shape species-specific bacterial clearance in Drosophilidae

**DOI:** 10.64898/2026.08.31.748369

**Authors:** Cong Li, Bing-Jun Wang, Junhui Peng, Nicolas Svetec, Li Zhao

## Abstract

Innate immunity has been dissected in exquisite detail in *Drosophila melanogaster*, yet how immune systems diversify between species remains largely unknown. Here we compare responses to bacterial infection across five drosophilid species spanning 60 million years of divergence, from *D. melanogaster* to *Scaptodrosophila lebanonensis*. Seven-day survival after Gram-negative infection ranges from 19% to 99% and tracks inversely with bacterial load in a phylogenetically corrected model, indicating that clearance rather than tolerance drives these differences. RNA sequencing of all five species after sterile wounding or infection reveals strongly species-biased transcriptional responses to the same pathogen, together with numerous uncharacterized lineage-restricted genes, including predicted antimicrobial peptides. We chemically synthesized candidate peptides and confirmed their activity in vitro: Athelas (CG43920), Mtkl and the *S. lebanonensis*-specific Athelas-like are active against bacteria and fungi, while Daisho2, previously described as antifungal, also kills Gram-positive and Gram-negative bacteria. De novo assembly and machine-learning prediction recover further candidate peptides from intronic and intergenic regions missed by current annotation. Rewiring of conserved genes and turnover of young, often unannotated effectors therefore act together to diversify antibacterial defense within a single insect family.

**One-sentence summary:** Five drosophilids over 60 million years defend against bacteria differently using lineage-specific young effectors and rewired conserved genes.

## Introduction

Living organisms inhabit complex and dynamic environments where exposure to microorganisms, including potential pathogens, is inevitable. To survive and maintain physiological homeostasis under such conditions, all organisms require effective immune defenses. In contrast to mammals, which possess both innate and adaptive immunity, fruit flies rely exclusively on innate immune responses, which include both humoral and cellular mechanisms^1–4^. The humoral immune response is primarily mediated through two major signalling pathways: the Toll pathway and the Immune deficiency (Imd) pathway, each specialized to target distinct classes of pathogens^5,6^. Upon activation, these pathways induce a rapid and robust expression burst of antimicrobial peptide (AMP) genes. Their expression typically increases by several hundred-fold, resulting in the secretion of multiple AMPs that synergistically neutralize and eliminate various microbial invaders^7–11^.

Immune-related genes typically evolve rapidly and exhibit strong footprints of positive selection due to the co-evolutionary arms race between host defense and pathogens^12–16^. Notably, despite their direct and critical roles in clearing microbial pathogens in the *Drosophila* immune system, AMPs evolve more slowly than many other immune-related genes, though with higher gene turnover rates^16–19^. Recent studies suggest that this slower rate might be due to balancing selection, which maintains genetic diversity by favoring different alleles under distinct ecological conditions^20–22^. These genetic variations are functionally relevant and associated with variation in pathogen susceptibility^23^.

Comparative studies across species offer a powerful approach to link genetic divergence with functional differences in immune responses^19,24^. Variations in pathogen pressures and ecological niches can drive species-specific evolution of immune gene regulation and effector activity^22^, resulting in divergent infection outcomes. Patterns of species-specific variation in individual AMP genes help explain observed differences in immune survival outcomes among species, and emphasize the potential for novel immune functions to arise through species- or lineage-specific genetic innovation. Indeed, lineage-specific genes often represent novel evolutionary origins of immune function. Evolutionarily new genes^25–28^, including those that originated de novo, may generate novel gene structures and novel functions^29,30^, thus expanding the functional gene space available to an organism, allowing adaptive specialization and increased phenotypic complexity. However, the extent to which variations in the composition and regulation of these immune-related genetic elements contribute to species-specific immune responses remains largely uncharacterized.

Beyond analysis limited to mature peptides and fully characterized proteins, recent studies have identified proteasome-mediated degradation products that function as encrypted proteins^31–33^ - bioactive fragments hidden within larger protein sequences. These discoveries support the hypothesis that some AMPs may be embedded within canonical proteins, representing an extensive yet largely unexplored reservoir of potential novel AMPs^34^.

The fruit fly *Drosophila* is an excellent model for studying innate immune responses. However, most immunological insights are derived from *D. melanogaster*, leaving it unclear how conserved these responses are across evolutionary time. To address this, we examined five fly species spanning a phylogenetic range from closely related to distantly related relatives of *D. melanogaster*: *D. melanogaster*, *D. simulans*, *D. ananassae*, *D. virilis*, and *Scaptodrosophila lebanonensis* to determine how immune responses vary across evolutionary distance.

To explore infection-induced variation in immune responses, we focused on two bacterial pathogens with ecological relevance to *Drosophila*. *Enterococcus faecalis* (Gram-positive) and *Providencia rettgeri* (Gram-negative) are opportunistic human pathogens frequently associated with nosocomial infections^35–39^. Both bacterial species are pathogenic to *D. melanogaster* flies and have been isolated from wild-caught flies^20,40^, and cause approximately 50% mortality by 7 days post-infection in *D. melanogaster*^41^. These characteristics make them ecologically relevant and representative natural pathogens for fruit flies, thus suitable models for testing interspecific variation in survival outcomes in this study. We characterize the variations in survival outcomes and immune responses across five fly species to *E. faecalis* or *P. rettgeri*, revealing that gene turnover and divergent regulation of conserved genes contribute to species-specific immune responses to the same pathogen challenge. We also identify evolutionarily new genes and unannotated genes with antimicrobial functions.

## Results

### Fly species differ markedly in survival following bacterial infection

To assess whether survival outcomes against two different pathogens vary across fruit fly species, we infected *D. melanogaster* (dmel), *D. simulans* (dsim), *D. ananassae* (dana), *D. virilis* (dvir), and *Scaptodrosophila lebanonensis* (sleb) with either *E. faecalis* or *P. rettgeri* (Figure 1A). Flies injected with sterile PBS solution served as a sterile wounding control, while a separate untreated group without any manipulation was used as a baseline control.

**Figure 1.**
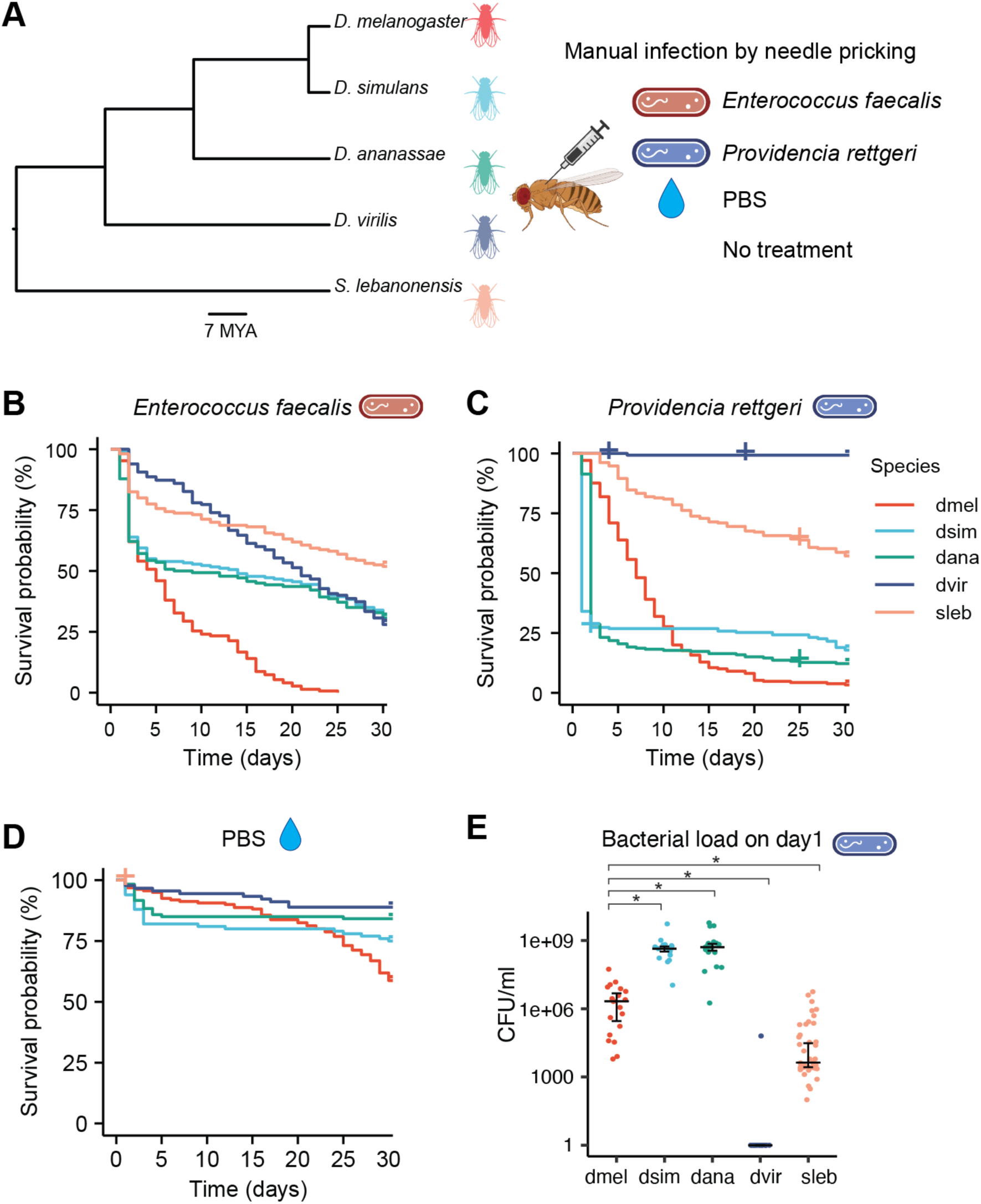
Species-specific survival performance correlates with bacterial load. A) Five fly species were manually infected with *E. faecalis*, *P. rettgeri*, or PBS as sterile wounding control; an untreated group was also included for RNA-seq analysis. B-C) Survival was monitored for 30 days following *E. faecalis* (B) and *P. rettgeri* (C) infection. Crosses indicate censored data. D) Survival curve after sterile wounding. E) Bacterial load measured one day after *P. rettgeri* infection (*, *p* < 0.05, one-sided Welch’s *t*-test with Holm correction).

To test whether our method successfully introduced bacterial infection and triggered immune activation in each species, we measured the expression of several AMP genes using quantitative RT-PCR one day post-infection (Figure S1C, S1D, S1F, S1H, S1J). Detected expression levels of AMP genes increased dramatically by up to ∼1,000-fold compared to sterile wound controls, consistent with previous studies^42–45^, confirming that our manual infection protocol effectively activated the immune responses.

With successful infection and immune activation, we observed striking differences in survival trajectories over time across the five species (Figure 1B, 1C). After *E. faecalis* infection, *D. melanogaster*, *D. simulans* and *D. ananassae* exhibited rapid mortality within the first two days, with survival rates at day 7 of 32.7%, 53.9% and 50.0% respectively (Table 1). In contrast, *D. virilis* and *S. lebanonensis* showed slower mortality over the 30-day period, particularly within the first 5 days, with day 7 survival rates of 86.0% and 73.8%, reflecting their high LT50 values. Following *P. rettgeri* infection, *D. simulans* and *D. ananassae* were acutely susceptible, with ∼75% mortality within two days and survival dropping to 26.8% and 18.6% by day 7. *D. melanogaster* and *S. lebanonensis* showed better early survival (48.1% and 83.3% at day 7) but experienced increased mortality over time. Remarkably, *D. virilis* displayed exceptional resistance, maintaining 99.3% survival at day 7, comparable to untreated controls. These results reveal strong species- and pathogen-specific variation in survival responses, suggesting that underlying genetic and transcriptional mechanisms differ substantially across species, even when exposed to the same pathogens.

**Table 1.** Survival outcomes after bacterial infection and comparisons with PBS sterile wound.

|  | *E. faecalis* |  | *P. rettgeri* |  | PBS |  | *E. faecalis* vs. PBS |  | *P. rettgeri* vs. PBS |  |
| --- | --- | --- | --- | --- | --- | --- | --- | --- | --- | --- |
| | Survival at 7 days | LT50 (day) | Survival at 7 days | LT50 (day) | Survival at 7 days | LT50 (day) | log rank test $\chi^2$ | p value | log rank test $\chi^2$ | p value |
| dmel | 32.7% | 5 | 48.1% | 7 | 91.3% | >30 | 270 | <2.2E-16 | 236 | <2.2E-16 |
| dsim | 53.9% | 14 | 26.8% | 2 | 82.0% | >30 | 44 | 3.0E-11 | 108 | <2.2E-16 |
| dana | 50.0% | 7 | 18.6% | 3 | 85.0% | >30 | 73 | <2.2E-16 | 168 | <2.2E-16 |
| dvir | 86.0% | 21 | 99.3% | >30 | 94.4% | >30 | 73 | <2.2E-16 | 14 | 1.4E-04 |
| sleb | 73.8% | >30 | 83.3% | >30 | 96.2% | >30 | 0.94 | 0.33 | 0 | 1 |

To test whether the observed differences in survival were primarily due to wounding from needle pricking rather than infection, we monitored the survival of PBS-treated (sterile wounding) flies under identical conditions. These flies showed significantly lower mortality compared to bacteria-infected groups (log-rank test: *p* < 0.001 for both *E. faecalis* vs PBS and *P. rettgeri* vs PBS in every species except *S. lebanonensis*), particularly during the first 2 days post-treatment, a period characterized by peak bacterial replication and acute immune activation^46^, indicating that mechanical injury alone did not account for most of the survival differences (Figure 1D). We also evaluated whether interspecific differences in body size influenced survival outcomes by infecting *D. virilis* and *S. lebanonensis*, two species with 2.00- and 1.24-fold larger body weight relative to *D. melanogaster*, with a 2-fold higher bacterial dose compared to the standard dose. The resulting survival curves closely matched those observed under standard dosing conditions, suggesting that body size did not significantly affect survival in our experimental setup (Figure S1A).

To investigate whether species-specific survival differences reflected variation in resistance (bacterial clearance) or immune tolerance^47–52^, we quantified bacterial load in individual flies one day after *P. rettgeri* infection. Bacterial colonization was confirmed in all five fly species, and bacterial load patterns corresponded with our observed survival outcomes. Specifically, *D. simulans* and *D. ananassae*, the two most susceptible species, carried bacterial loads more than two orders of magnitude higher than *D. melanogaster* (Figure 1E). *S. lebanonensis*, which showed intermediate survival after 7 days, had a >1 magnitude lower bacterial load than *D. melanogaster*, though not significant (Figure 1E). Remarkably, *D. virilis*, which showed a near 100% survival rate, carried bacterial loads near baseline levels and close to the detection limit - over three orders of magnitude lower than *D. melanogaster* (Figure 1E). A Bayesian analysis considering phylogenetic information revealed a negative correlation (mean correlation coefficient ρ = - 0.875) between bacterial load at day 1 and survival at day 3 across species (Figure S1B), indicating that bacterial burden strongly predicts susceptibility to infection. Since all species were exposed to the same initial bacterial dose relative to body size, the differences in survival outcomes indicated that differences in bacterial clearance, rather than tolerance, injury susceptibility, or body size, are the primary drivers of survival variation. Together, these findings demonstrate that fruit fly species differ markedly in their ability to survive bacterial infection, primarily due to differences in bacterial clearance.

### Transcriptome profiling reveals high divergence of gene expression regulation in immune responses

To comprehensively profile immune responses against the two pathogens and explore mechanisms underlying survival differences, we conducted bulk RNA sequencing on samples collected one day post-infection, a time point at which AMP gene expression peaks^46^. We generated three biological replicates per treatment group, including untreated, PBS (sterile wounded), and bacteria-infected conditions across two bacteria and five fruit fly species, resulting in 90 total libraries. Robust transcriptional immune activation was observed after bacterial infection in *D. melanogaster* (Figure 2A, 2B) and the other four species as well (Figure S1C-S1K). In *D. melanogaster*, 586 and 186 genes were significantly upregulated following *E. faecalis* and *P. rettgeri* bacterial infection, respectively, while 48 and 31 genes were significantly downregulated. Gene Ontology (GO) enrichment analysis revealed that the top enriched terms among upregulated genes were immune-related (Figure 2C, 2D, Figure S2C, S2D), including several effector genes that have been implicated in antimicrobial roles, such as *Dro*, *Dpt*, *Cec*^44^ (Table S1). Among them, some genes were also induced by sterile wounding (Table S2). These include members of the *Bomanin* and *Turandot* gene families, which are known to be activated by diverse stress conditions^53–56^, including immune responses^23,57–62^.

**Figure 2.**
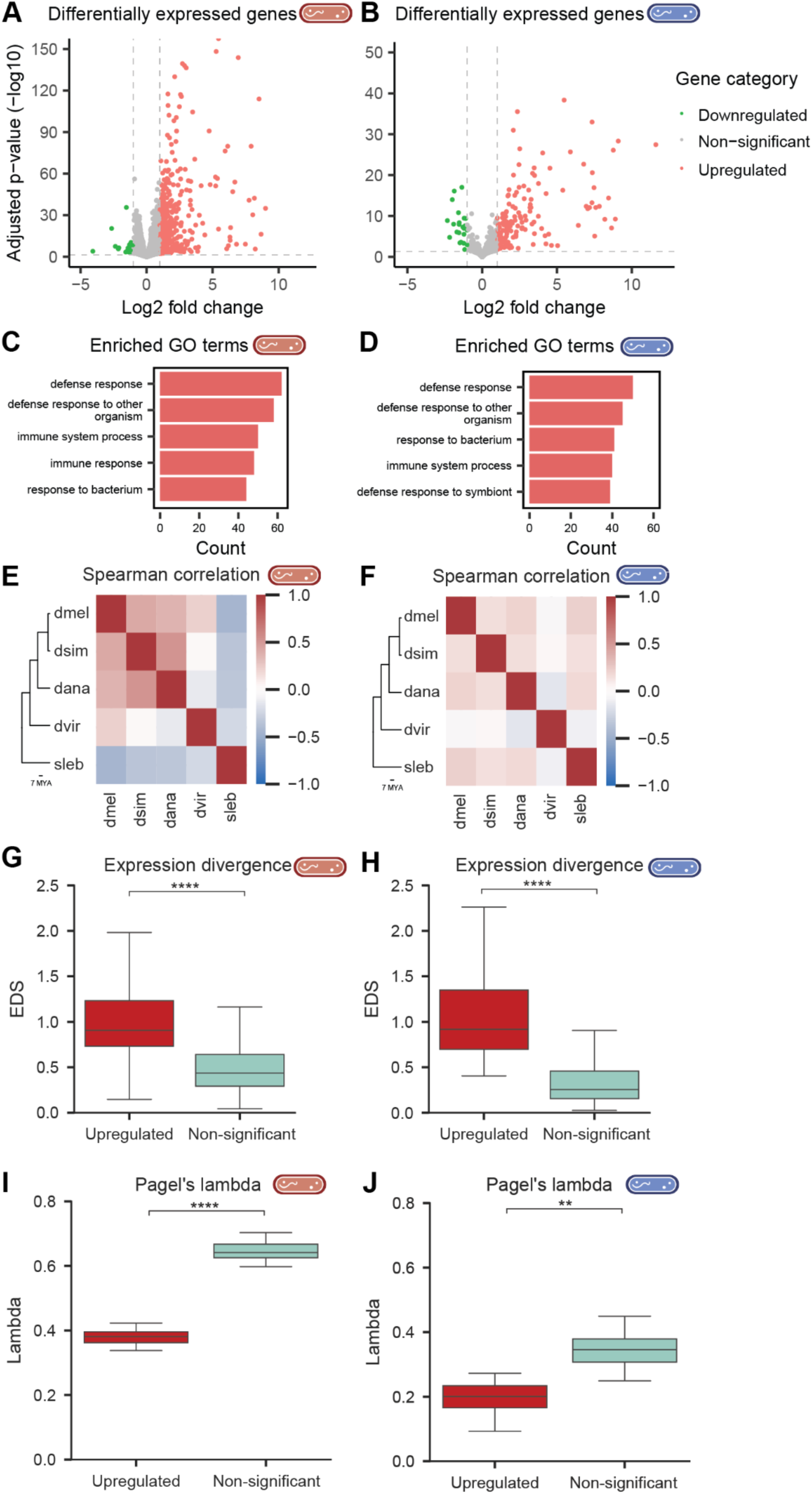
Gene expression profiles in *D. melanogaster* and divergence across species following bacterial infection. A-B) Volcano plots showing differentially expressed genes in *D. melanogaster* after infection with *E. faecalis* (A) and *P. rettgeri* (B). C-D) GO analysis shows enriched terms related to immune response. E-F) Heatmaps of genome-wide Spearman correlations in log_2_ fold-change across species for *E. faecalis* (E) and *P. rettgeri* (F) infections. Positive values indicate concordant responses; negative values indicate divergence. G-H) Upregulated genes exhibit significantly higher Expression Divergence Scores (EDS) than non-significant genes following *E. faecalis* (G) and *P. rettgeri* (H) infections. EDS was calculated as the mean of all pairwise absolute differences in log_2_ fold-change between species, with higher values reflecting greater interspecies variability in infection-induced responses. Upregulated genes are defined as those induced in at least one species in a bacteria-infected group compared to their correlated untreated group; non-significant genes show no induction in any species. ****, *p* < 0.0001, one-sided Welch’s *t*-test. I-J) Upregulated genes show significantly lower lambda values on the log_2_ fold-change following *E. faecalis* (I) and *P. rettgeri* (J), indicating weaker phylogenetic signals ****, *p* < 0.0001, **, *p* < 0.01, Mann-Whitney U test.

To identify genes specifically induced by bacterial infection independent of wounding effects, we compared gene expression between bacteria-infected and sterile-wounded groups. This comparison revealed 60 genes (in *E. faecalis*) and 114 genes (in *P. rettgeri*) that were significantly upregulated in the bacterial-infected condition relative to sterile wounding. Notably, several genes such as *DptA* and *CecA1* exhibited over 1,000-fold higher expression in bacteria-infected flies, consistent with our qPCR results (Table S2 and Figure S1C). Many of these infection-induced genes were associated with pathways triggered by microbial components such as lipopolysaccharides^63,64^, further underscoring their specificity to pathogenic challenge. Across the four non-*melanogaster* species, both *E. faecalis* and *P. rettgeri* infections elicited substantial transcriptional responses, with many genes upregulated and a smaller number of genes downregulated in most cases (Figure S1D-K, Table S1).

To compare expression patterns of immune-responsive genes across species, we identified orthologous genes using OrthoMCL^65^, a genome-scale algorithm for detecting orthologous protein sequences. We identified 9,296 one-to-one orthologous gene groups in which a single-copy gene was present in all five species (Figure S2A, File S1). Principal component analysis (PCA) based on the top 500 most variable genes revealed strong species-specific clustering (Figure S2E, S2F), suggesting evolutionary divergence of infection-induced transcriptional responses across fly species. Notably, *D. simulans* and *D. ananassae*, the two species that exhibited the highest early mortality following both *E. faecalis* and *P. rettgeri* infection (Figure 1B, 1C), clustered closely together and distinctly from the other three species. This clustering may reflect transcriptional divergence in immune responses that contributes to different survival outcomes, or alternatively, a shared acute infection state resulting from higher susceptibility. PCA loadings positively correlated with PC1 were enriched for GO terms such as “defense response” and “immune response”, further supporting that variation in immune-related gene expression is associated with species-specific infection responses (Table S3).

To compare expression divergence across species, we calculated Spearman correlations of log_2_ fold-changes in all one-to-one orthologous genes in response to bacterial infection relative to the untreated condition (Figure 2E, 2F). *S. lebanonensis* showed low correlation with other species in *E. faecalis* infection. *D. virilis* showed consistently low correlation with other species in both infection conditions, especially with *D. simulans* and *D. ananassae*, which had the worst survival performance.

To quantitatively assess divergence in regulation of genes across species, we computed Expression Divergence Scores (EDS) for each one-to-one orthologous gene (see Methods). We found that genes that were upregulated in at least one species exhibited significantly higher EDS values than genes that were not significantly differentially expressed in any species (Figure 2G, 2H, Table S4), indicating that infection-responsive genes tend to show greater expression divergence across species. This divergence in expressions of upregulated genes may contribute to the variance of survival outcomes observed above.

To identify candidate genes that may contribute to species-specific divergence in immune responses and potentially influence survival outcomes, we focused on *D. virilis*, the species with the highest survival rate. Specifically, we selected genes that were upregulated in *D. virilis* and showed both high expression levels (baseMean > 1,000) and strong expression divergence across species (EDS > 2.0; Table S4). These thresholds were chosen to prioritize genes with both robust activation and high species-specificity, which are more likely to underlie functional differences in infection outcomes. Among those responding to *E. faecalis* and/or *P. rettgeri*, the top genes were *Dso2*, *DptB*, *BomBc3*, *Ilp8* (Insulin-like peptide 8), *PGRP-SD, Def* and *Obp99b* (odorant-binding protein 99b). Several of these genes, such as *DptB* and *Def*, are well-known antimicrobial peptides^20,22,23,66–68^, while others like *Ilp8* - a stress-responsive peptide involved in systemic growth coordination and development - suggest broader integration with physiological processes such as insulin signalling during immune responses^69,70^. Notably, *Obp99b* was specifically upregulated in *D. virilis* and displayed both high expression and high EDS, suggesting a potential role in chemosensory-driven modulation in response to infection. Among the seven genes, *DptB, Def* and *Obp99b* showed the highest induction in *D. virilis* compared to the other four species. These genes, which show high variance in expression after infection, represent candidate effectors and regulatory targets that may underlie the enhanced immune performance of *D. virilis*, providing a potential mechanistic link between species-specific transcriptional divergence and improved survival outcomes.

To further investigate whether the evolution of immune responses contributes to differences in bacterial infection resistance across the five species, we mapped the evolutionary patterns of infection-induced gene expression and assessed phylogenetic signals using Pagel’s lambda^71–73^. This metric quantifies the extent to which variation in a trait can be explained by the phylogeny. We compared lambda values between upregulated genes and genes that were not significantly upregulated, and found that upregulated genes exhibited significantly lower lambda values (Figure 2I, 2J). This suggests that the divergence in expression of immune-responsive genes is less constrained by the phylogenetic relatedness, potentially reflecting species-specific evolution of gene regulation^74^.

Together, these findings revealed substantial transcriptional differences in infection-responsive genes after infection across species, indicating that evolutionary divergence in gene regulation may contribute to phenotypic variation such as differences in survival.

### Lineage-biased immune responses arise from divergent regulation of conserved genes

To further investigate the molecular basis of species-biased immune responses, we examined how both conserved and lineage-specific genes contribute to the species-specific immune responses. The 9,296 one-to-one conserved orthologs generally exhibited similar expression profiles after *E. faecalis* and *P. rettgeri* infections (Figure S3A-S3D and File S1). However, their infection-induced responses showed only moderate overlap across species (Figure 3A and S3E).

**Figure 3.**
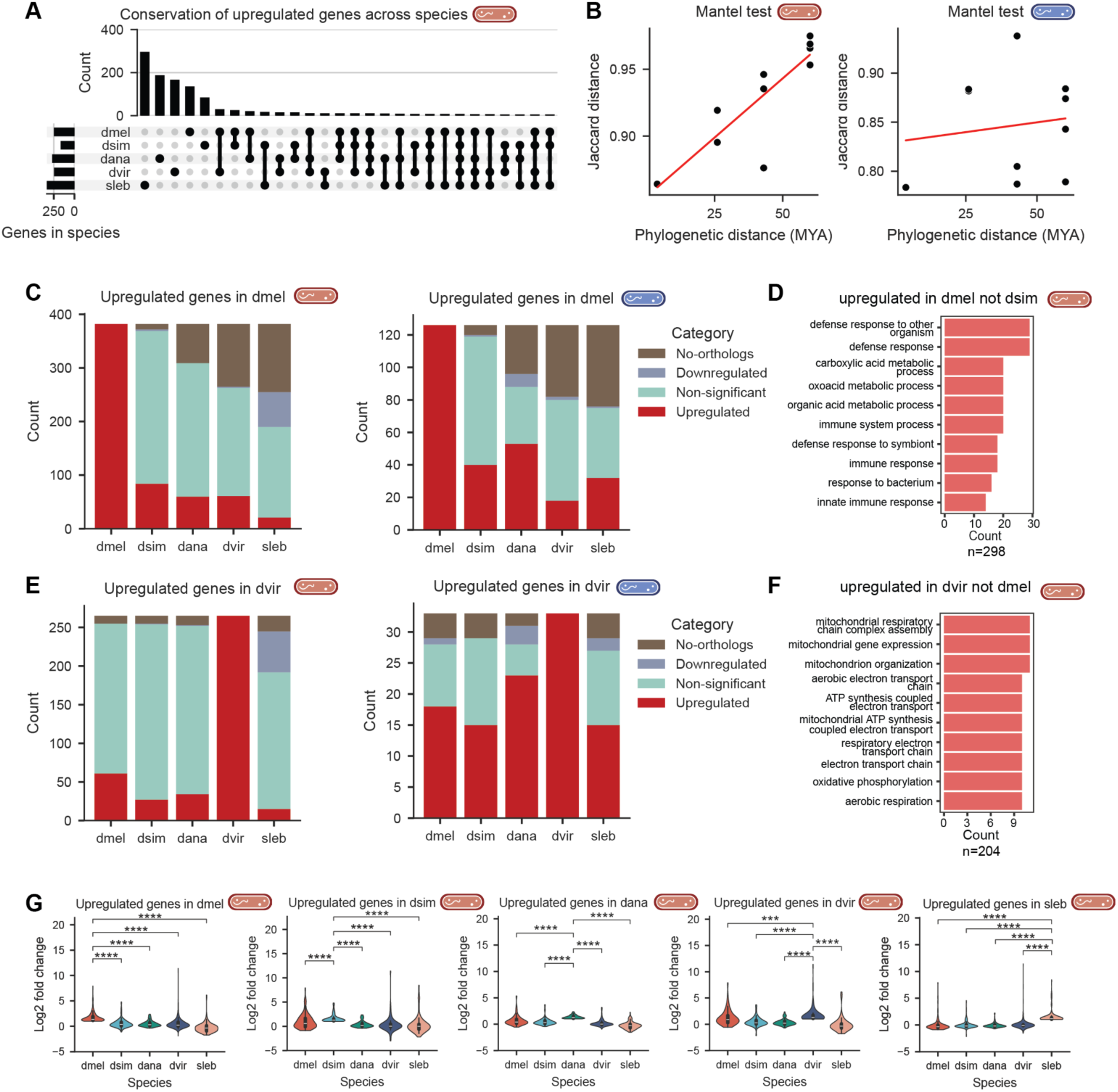
Shared gene regulation and species-specific immune responses. A) The UpSet plot illustrates, for each orthologous gene group, which species exhibit significant upregulation following *E. faecalis* infection. Although overall expression levels are broadly conserved, upregulation in response to infection is not consistently shared across species. B) Mantel test results show the relationship between immune-response similarity and phylogenetic distance across species. Following *E. faecalis* infection, similarities in gene upregulation were positively correlated with phylogenetic distance (Mantel *r* = 0.849, *p* = 0.0180). In contrast, the correlation was weaker and not significant following *P. rettgeri* infection (Mantel *r* = 0.142, *p* = 0.7710). C) Among orthologous genes upregulated in *D. melanogaster*, some lack identifiable counterparts in other species, while others show divergent regulatory responses. These plots show, for each *D. melanogaster* upregulated gene, the direction of regulation (upregulated, downregulated, or not significant) in each species where an ortholog is present. Moreover, the proportion of *D. melanogaster* immune genes lacking orthologs increased with phylogenetic distance (e.g., 33% absent in *S. lebanonensis*). D) Genes upregulated in *D. melanogaster* but not in *D. simulans* are enriched for metabolic processes. E) Similarly, many genes upregulated in *D. virilis* are either absent or differently regulated in other species. These plots show, for each *D. virilis* upregulated gene, the direction of regulation in each species where an ortholog is present. F) Genes upregulated in *D. virilis* but not in *D. melanogaster* are enriched for respiratory-related processes. G) Genes upregulated in one focal species after *E. faecalis* infection show significantly higher fold changes in that species compared to their orthologs in other species (*, *p* < 0.05; **, *p* < 0.01; ***, *p* < 0.001; ****, *p* < 0.0001; Mann–Whitney U test with FDR-Benjamini Hochberg correction).

To investigate how regulation of immune-induced genes varies across species, we analyzed 1,005 one-to-one orthologous gene groups in which at least one species showed upregulation following *E. faecalis* infection. The majority of upregulation patterns were species-specific, with the highest number of genes observed in *S. lebanonensis* (292 genes), followed by *D. ananassae* (183), *D. virilis* (162), *D. melanogaster* (131), and *D. simulans* (80). A similar pattern was observed following *P. rettgeri* infection, where among 390 upregulated orthologous groups, most were restricted to one or two species: 214 in *D. ananassae*, 42 in *S. lebanonensis*, 26 in both *D. ananassae* and *S. lebanonensis*, and 23 in *D. melanogaster*. Only three genes were consistently upregulated in all five species after *E. faecalis* infection, including the signalling gene *PGRP-SB1* and the effector gene *DptB*. After *P. rettgeri*, nine genes were commonly upregulated across species, including signalling genes *PGRP-SA*, *PGRP-SB1*, *PGRP-SD*, and *PGRP-LF*, as well as effector genes *DptB*, *Def*, *AttD*, and *IM33*. Beyond genes upregulated in a single species, some genes were co-regulated across multiple species. For example, 12 genes were shared across four species after *E. faecalis* infection, including *BomBc3*, *BomT1*, *Dso1*, *Dso2*; 8 genes were shared after *P. rettgeri* infection, including *PGRP-LB*, *BomBc3*, *BomT1*, and *edin*. These results suggest that while immune-induced gene expression is largely species-specific, a small set of key immune signalling and effector genes show conserved upregulation across species.

To validate this pattern, we focused on the one-to-one orthologs upregulated in *D. melanogaster* (382 and 126 orthologous genes following *E. faecalis* and *P. rettgeri* infection, respectively; Figure S1, File S1). Only a portion of these were also co-upregulated in other species (Figure 3C; for *E. faecalis*: 84 in *D. simulans*, 60 in *D. ananassae*, 61 in *D. virilis*, 21 in *S. lebanonensis*; for *P. rettgeri*: 40 in *D. simulans*, 53 in *D. ananassae*, 18 in *D. virilis*, 32 in *S. lebanonensis*). Many showed no significant expression changes (44.2% - 74.6% in the four species) or were even downregulated (e.g., 65 genes in *S. lebanonensis* after *E. faecalis* infection). Following *P. rettgeri* infection, the non-*D. melanogaster* species also displayed more genes with no change or no identifiable orthologs than genes with upregulation (combined non-significant and no-ortholog categories: 51.6% - 84.1% across species).

To formally test whether closely related species share more upregulated genes, we performed a Mantel test. The results revealed that similarity in gene upregulation was positively associated with phylogenetic distance following *E. faecalis* infection (Mantel *r* = 0.849, *p* = 0.018; Figure 3B), whereas no significant relationship was observed for *P. rettgeri* (Mantel *r* = 0.142, *p* = 0.771; Figure 3B). Together, these findings suggest that regulatory divergence among conserved genes contributes to species-specific immune responses, and that immune response regulatory divergence may itself be pathogen specific. Additionally, gene turnover, particularly in more distantly related species, may further contribute to differences in infection-induced responses.

GO enrichment analysis further supported this interpretation. Genes upregulated in both *D. melanogaster* and *D. simulans* following *E. faecalis* infection were enriched for immune-related functions, including 69 genes associated with the GO term “Signal” and 7 AMP genes (Figure S3F). In contrast, genes upregulated in *D. melanogaster* but not in *D. simulans* were enriched for various metabolic processes in addition to immune responses (Figure 3D). Among these 298 genes, 187 genes were associated with the term “Signal”, but only 2 AMP genes - *AttD* and *BomS4* - were included. This suggested that signalling components of the immune response may evolve more rapidly than effector genes such as AMPs, which is consistent with previous findings^17–19^. Overall, this pattern indicates that conserved infection-responsive genes are more likely to encode core immune components, e.g. effectors such as AMPs, whereas genes specifically induced in *D. melanogaster* may reflect species-specific physiological programs or secondary responses to infection, such as stress-associated metabolic changes^75–80^ or non-canonical metabolic functions of pattern-recognition proteins such as cGAS/cGLR-family enzymes^81–83^.

*D. virilis*, the species with the highest survival rate, exemplified this divergence (Figure 3E). Specifically, 265 genes were upregulated after *E. faecalis* infection, the majority of which were not significantly induced in other species (66.8% - 85.7%). Following *P. rettgeri* infection, fewer genes were upregulated in *D. virilis*, and approximately half were shared with other species (Figure 3E). Notably, genes specifically induced in *D. virilis* but not in *D. melanogaster* were enriched for GO terms related to mitochondrial and respiratory functions (Figure 3F), suggesting that *D. virilis* may have distinct transcriptional programs targeting mitochondrial activity during infection. Many of these genes are associated with transit peptides - N-terminal sequences that direct newly synthesized proteins to organelles such as mitochondria. This enrichment points to a potential role for enhanced mitochondrial targeting, signalling and function in *D. virilis*, which may underlie its increased resilience to infection^84–86^.

Consistent with the limited overlap of significantly upregulated genes, genes significantly upregulated in each species after *E. faecalis* infection displayed significantly greater fold changes in the focal species relative to their orthologs in other species (> 2-fold difference; Figure 3G). A similar, though less pronounced, pattern emerged following *P. rettgeri* infection (Figure S3G). These findings further support the notion that regulatory divergence among conserved genes plays an important role in shaping species-specific immune responses.

Together, these findings reveal that despite a conserved core of immune genes including signalling and effector genes, infection-induced transcriptional responses are often species-specific, suggesting distinct adaptive strategies potentially involving metabolic or physiological shifts. Regulatory divergence is common even among highly conserved one-to-one orthologous genes, suggesting rapid evolution of immune gene regulation shaped by species- or lineage-specific ecological pressures. In addition, gene turnover contributes further to the diversification of immune responses across the fruit fly phylogeny.

### Between-species differential expression analysis identifies genes underlying species-specific immune response and survival outcome

To uncover genes with divergent expression patterns that may contribute to species-specific immune responses and survival differences following bacterial infection, we compared gene expression profiles across species under bacteria-infected conditions. Based on survival data, we defined two species groups: a susceptible group (*D. melanogaster*, *D. simulans, D. ananassae*), and a resistant group (*D. virilis*, *S. lebanonensis*). Using two complementary approaches - a fold-change threshold applied to within-species induction, and a direct between-species differential expression test (see Methods) - we screened one-to-one orthologs to identify genes specifically upregulated in the resistant group. This analysis yielded 77 genes for *E. faecalis* and 23 genes for *P. rettgeri* (Table S5). These genes were enriched for GO terms related to secretion, immunity and mitochondrial activity, and included immune-related genes (*Defensin*, *PGRP-SB1*, *PGRP-SD*, *Diedel*, *CG6421*) as well as olfaction-related genes (*Obp99b*, *Obp44a*). For example, *Defensin* expression after *E. faecalis* infection closely mirrored the observed survival patterns - remaining low in the susceptible group and strongly upregulated in the resistant group (Figure 4E). *Def* encodes an AMP in *D. melanogaster* that is highly effective against Gram-negative Providencia, but with relatively low expression^66,67^. Its robust induction in resistant species suggests a species-specific regulatory mechanism that may contribute to enhanced survival, in addition to possible effects of sequence divergence, as *Defensin* orthologs are present in all five species.

**Figure 4.**
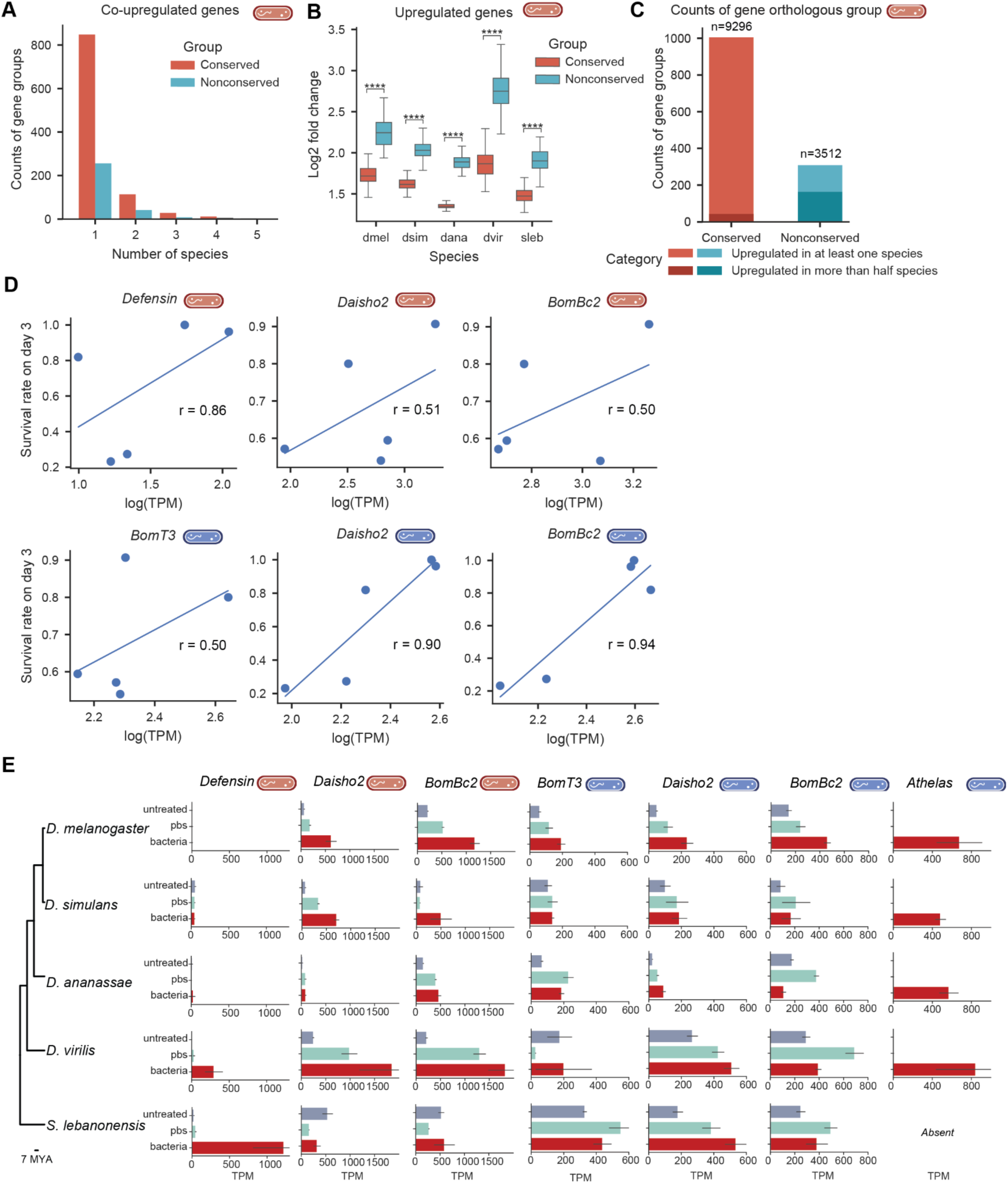
Small, rapidly evolving genes contribute to immune responses following bacterial infection. A) Most orthologous gene groups show upregulation in only one single species after *E. faecalis* infection, suggesting species-specific immune responses. B) Non-conserved genes exhibit significantly stronger induction upon infection (****, *p* < 0.0001; *p* values were calculated by one-sided Welch’s *t*-test) Gene lists were selected using simulated annealing to match expression distributions. C) Non-conserved genes are more likely to be co-regulated across species (induction ratio: 0.528 vs. 0.043*; p < 0.001,* Fisher’s exact test). D) Expression of *Defensin*, *Daisho2*, and *BomBc2* is positively associated with survival rate on day 3 after *E. faecalis* infection (Pearson correlation, r values shown in the plots; correlation mean = 0.85, 0.55, and 0.40; β₁ mean = 0.15, 0.15, and 0.20, respectively), while expression of *Daisho2*, *BomBc2* and *BomT3* is positively associated with survival rate on day 3 after *P. rettgeri* infection (Pearson correlation, r values shown in the plots; correlation mean = 0.90, 0.96, and 0.67; β₁ mean = 1.09, 1.19, and 1.28, respectively). E) Expressions of highly correlated genes after *E. faecalis* infection or *P. rettgeri* infection in the species where they are present. Bars represent mean TPM with standard error.

We also examined the correlation between gene expression and survival across species with 15 single-copy candidate or known AMP genes. Expression of the top three genes - *Defensin*, *Daisho2*, *BomBc2* - was positively associated with survival following *E. faecalis* infection (Figure 4D-E), and expression of *Daisho2*, *BomBc2,* and *BomT3* was positively associated with survival following *P. rettgeri* infection (Figure 4D-E). These results highlight a set of candidate genes whose divergent expression may drive species-specific immune strategies and contribute to differential survival outcomes following infection.

### Small, rapidly evolving genes contribute to immune responses to bacterial infection

Our results suggest that conserved genes contribute to species-specific responses to a shared bacterial challenge across five species by regulating a common set of genes in a species-dependent manner. Although conserved genes constitute the majority of protein-coding genes in *D. melanogaster* (10,440 genes; 78.8%; Figure S2A), a substantial number of non-conserved genes (4,165), which are absent from one or more of the five species, also participate in immune responses. Most orthologous gene groups, both conserved (84.4%) and non-conserved (82.8%), were upregulated in only a single species following *E. faecalis* infection (Figure 4A). Nonetheless, non-conserved genes tended to be more strongly induced upon infection (Figure 4B) and showed a higher frequency of upregulation across multiple species (Figure 4C). Together, these patterns indicate that evolutionarily dynamic (non-conserved) genes are more strongly activated in response to bacterial infection.

Among both conserved and non-conserved genes, those encoding short peptides - typically under 200 amino acids - are of particular interest, as many exhibit features consistent with antimicrobial function, such as the presence of N-terminal signal peptides. Notably, the expression levels of several such genes were strongly correlated with host survival following bacterial infection (Figure 4D). Building on our RNA-seq analysis, we identified 32 and 19 small, rapidly evolving genes that were strongly induced by *E. faecalis* and *P. rettgeri* bacterial infection, respectively (Table S6). To estimate their evolutionary origins and trajectories, we mapped their phylogenetic distribution, referring to a previously established phylogenetic framework for *D. melanogaster* genes^30^. Single-cell RNA-seq data revealed that 16 out of 32 and 11 out of 19 genes were highly expressed in immune-relevant tissues, including the fat body and hemocytes^87–93^, in *E. faecalis* and *P. rettgeri* datasets respectively, further supporting their potential roles as functional components in immunity.

For example, *Daisho2* (FBgn0067905), previously reported to have antifungal activity^94^, was robustly upregulated following bacterial infection in all five species analyzed, and was highly expressed even in untreated *D. virilis* and *S. lebanonensis* flies (Figure 4E). To evaluate the antimicrobial properties of these peptides, we chemically synthesized the encoded peptide and tested its activity *in vitro* using minimum inhibitory concentration (MIC) assays^95,96^ against a panel of bacterial and fungal pathogens relevant to insect and human health. Daisho2 inhibited the growth of nearly all tested bacteria, in addition to its previously reported antifungal activity, indicating that it functions as a broad-spectrum antimicrobial peptide. Microscopic examination revealed that bacterial cultures treated with Daisho2 became visibly clear relative to the peptide-free control (Fig S4E), consistent with bacterial cell lysis.

Besides *Daisho2*, we focused on two genes that are predicted to encode antimicrobial peptides but whose functions have never been examined experimentally: *Mtk-like* (*Mtkl*) and *CG43920*. We named *CG43920* as *Athelas*, after a fictional healing herb, reflecting its putative role in defense. *Mtkl* is named for its shared Antimicrobial10 domain^41,97^ with the well-characterized AMP gene *Metchnikowin* (*Mtk*)^98,99^. The *Mtkl* gene could be identified in most Drosophilidae species except *S. lebanonensis* (Figure 5A), and most orthologs of *Mtkl* encode a polyproline motif as well as a conserved PSPFNP motif at the C-terminus (Figure S4A). In the syntenic region of *S. lebanonensis*, we could not identify the gene, but recovered an unannotated transcript showing very high expression after both *E. faecalis* and *P. rettgeri* infections (Figure S4D). The transcript could potentially encode a peptide containing a signal peptide (Figure S4B). Different from *Mtkl*, the encoded peptide lacks a polyproline motif. However, it is most similar to *Athelas* when comparing it with all annotated genes in *D. melanogaster*; we therefore denote it as *Athelas-like* (Figure S4B). In the orthologous region of *Chymomyza caudatula* and *C. procnemis*, ORFs encoding peptides similar to *Athelas-like* were identified alongside the *Mtkl* orthologs (Figure 5A), while absent in all Drosophila species. By contrast, the orthologs of *CG43920* are conserved across all species in the genus *Drosophila*, but absent from the syntenic region in the other three Drosophilinae species (including *S. lebanonensis*, *C. caudatula*, and *C. procnemis*) (Figure 5A). Similar to the upregulation of *Athelas-like* after infections, *Athelas* showed higher expression after infections across all four Drosophila species in our experiments (Figure 4E). The protein sequences encoded by the genes in the orthologous regions of *Athelas-like* and those of *Athelas* are similar and have a conserved TPPFNP motif (Figure S4B).

**Figure 5.**
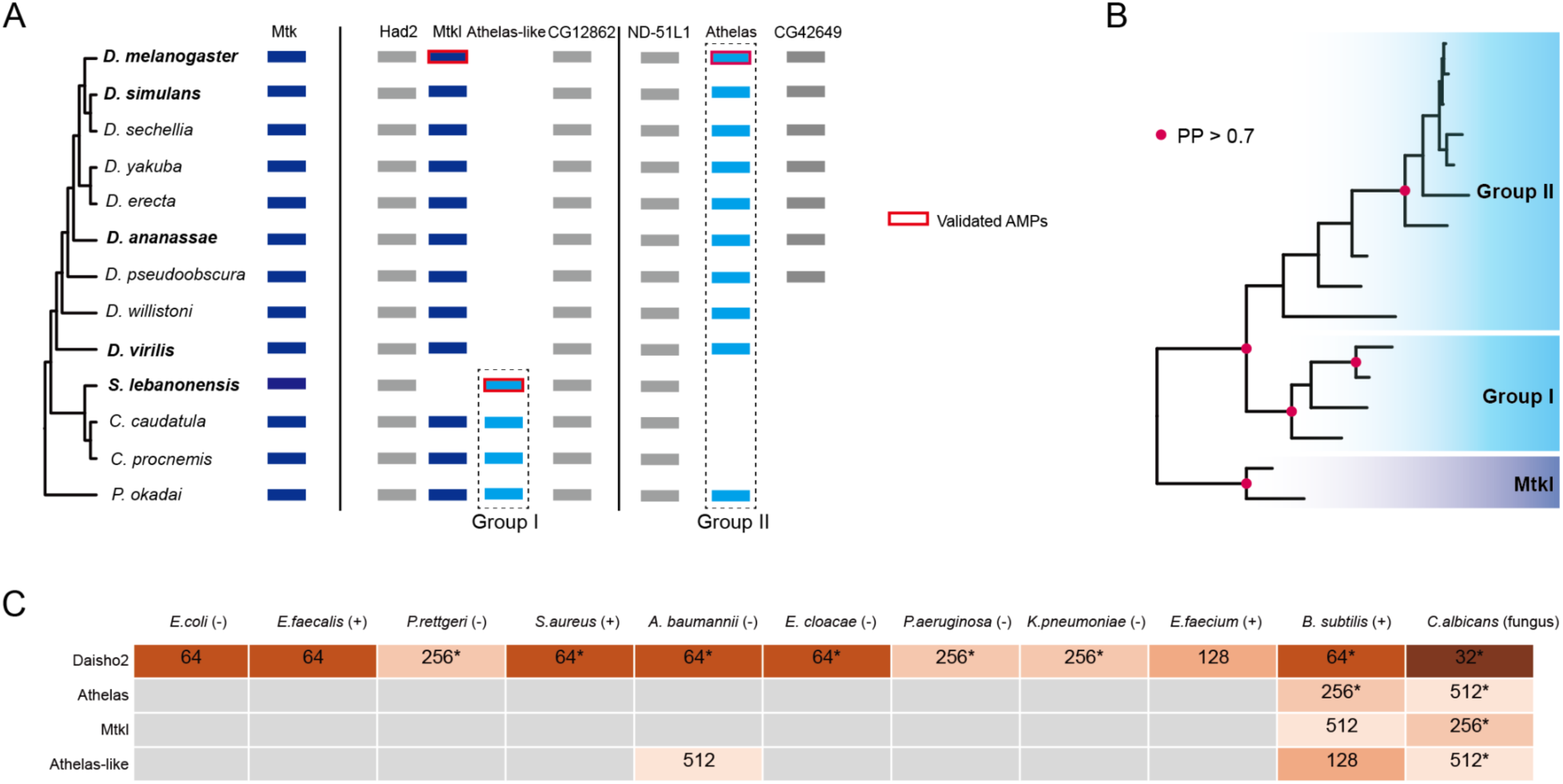
Evolutionary analysis and functional examination of *Mtkl*, *Mtk*, *Athelas, Athelas-like* genes. A) *Mtk* is conserved across all species examined, whereas *Mtkl* was found in most species except *S. lebanonensis*. Homologs of *Athelas* are shown in light blue and classified into Group I and Group II based on their genomic positions. Group I genes are positioned adjacent to *Mtkl* and were identified only in non-*Drosophila* species, while Group II genes were found in *Drosophila* species and *P. okadai*. Flanking genes used to define orthologous regions are shown in gray. Genomic segments that are not contiguous are separated by vertical lines. Genes with experimentally verified antimicrobial activity are represented by solid red lines. Species for which survival assays and infection-induced transcriptome profiling were performed are highlighted in bold in the phylogenetic tree. B) Phylogenetic analysis of *Athelas* homologs. *Mtkl* from *D. melanogaster* and *P. okadai* were used as outgroups. Genes from Group I and Group II each clustered according to their respective groups, forming sister clades. Nodes with posterior probabilities greater than 0.7 are highlighted with red dots. C) In vitro assays demonstrated antimicrobial activities of the corresponding peptides on bacteria and fungi. Gram-negative and Gram-positive bacteria are indicated as (-) and (+), respectively. Values denote the minimum concentration (mg/mL) required to inhibit bacterial growth; darker shades indicate lower MIC values. Asterisks mark cases of incomplete inhibition, characterized by residual turbidity or white floc formation.

Therefore, they may be homologs and we named them Group I and Group II, respectively (Figure 5A). In one species from Steganinae, *Phortica okadai*, which forms a sister group to all other Drosophilinae species, both Group I and Group II genes could be identified (Figure 5A). It is possible that the common ancestor of Drosophilidae possessed *Mtkl* as well as both Group I and Group II homologs of *Athelas*, and Group I was subsequently lost in Drosophila, whereas Group II was lost in some other Drosophilinae species; *Mtkl* was lost in *S. lebanonensis*. Phylogenetic analysis of the mature region sequences showed that Group I and Group II genes each clustered together according to their respective groups, forming sister clades, further supporting our inference of gene evolution based on synteny (Figure 5B). Based on the sequence similarity among *Mtk*, *Mtkl*, and *Athelas* - including a shared consensus motif (PF/YNP) at the C-terminus - they may belong to the same gene family (Figure S4A-C).

Functional studies showed that Athelas and Mtkl from *D. melanogaster* inhibited the growth of *B. subtilis* bacterium and *C. albicans* fungus, and Athelas-like in *S. lebanonensis* inhibited *A. baumannii* bacterium in addition to *B. subtilis* and *C. albicans* (Figure 5C). CG42649, encoded by a gene adjacent to *Athelas* (Figure 5A) and also containing a signal peptide, was not tested for antimicrobial activity, as its lack of induction upon infection suggested limited involvement in immune defense. Together, these results suggest that short, rapidly evolving genes - many of which lack annotation or conservation across species - can encode functional antimicrobial peptides.

### Unannotated open reading frames potentially encode antimicrobial peptides

Unannotated open reading frames (ORFs) represent a promising yet underexplored source of antimicrobial peptides (AMPs), especially in non-model species. In *D. melanogaster*, where gene annotation is relatively complete, most unannotated sequences (∼84%) are composed of repetitive elements. In contrast, a substantial fraction of unannotated genomic sequences in non-*melanogaster* species - *D. simulans* (33%)*, D. ananassae* (75%)*, D. virilis* (71%)*, S. lebanonensis* (73%) - cannot be explained solely by simple repeats. A higher proportion of newly assembled transcripts in these species (41% - 59%) also contain less than 5% repetitive sequence (Figure 6A, B).

**Figure 6.**
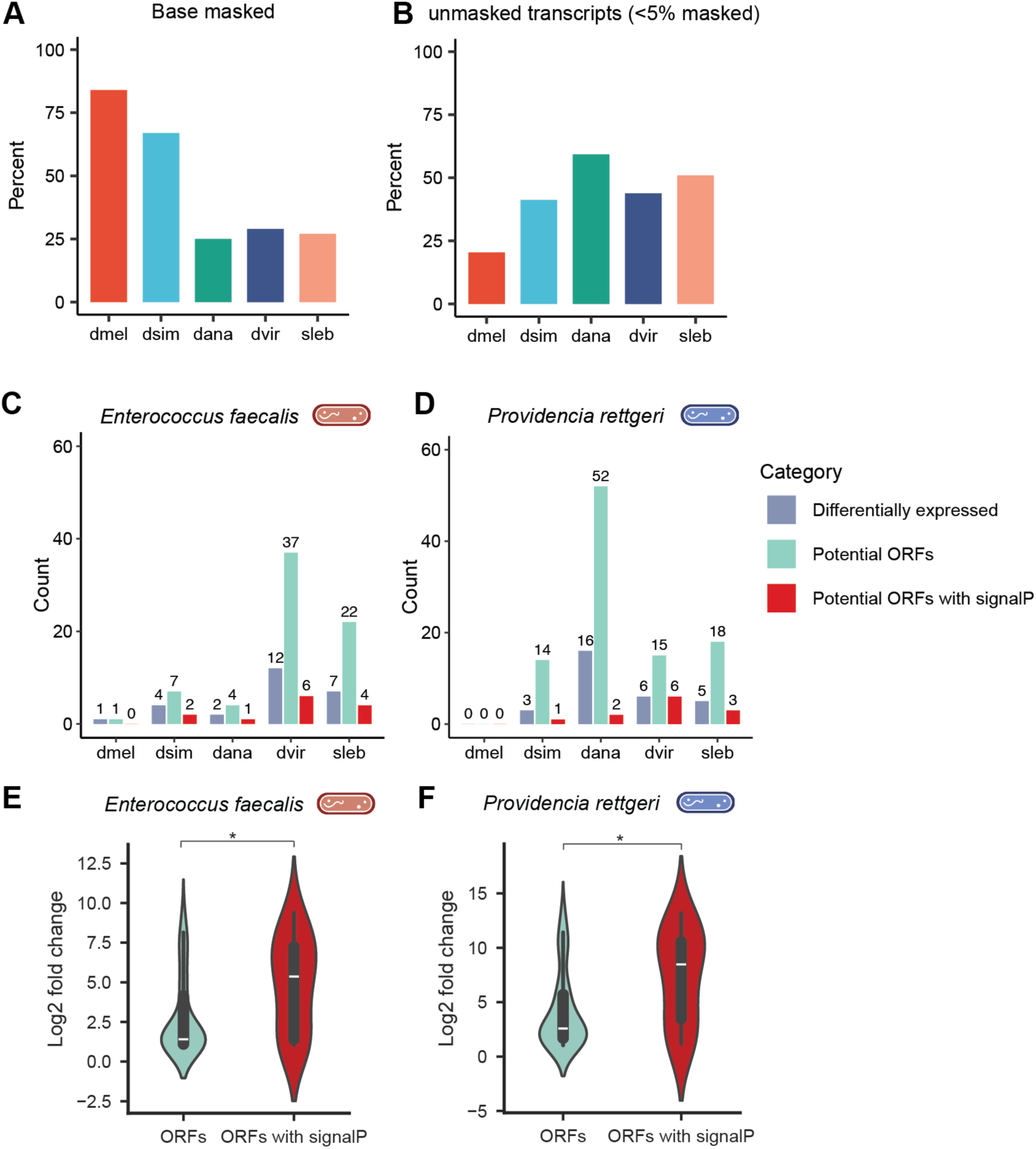
Evolutionarily new genes contribute to immune responses to bacterial infection. A) In *D. melanogaster*, most unannotated sequences are repetitive, whereas a large proportion in non- *melanogaster* species are non-repetitive. B) Many newly assembled transcripts in non-*melanogaster* species contain less than 5% repetitive content. C-D) From these unannotated transcripts, upregulated ones were identified, and open reading frames (ORFs) containing signal peptides were selected as candidates for encoding antimicrobial peptides (AMPs). E-F) ORFs with predicted signal peptides show significantly higher induction upon infection compared to the broader set of ORFs (*, *p* < 0.05; one-sided Welch’s *t*-test).

Many unannotated transcripts not captured in reference annotations or annotated non-coding regions contain potential ORFs, suggesting they may encode functional peptides^100,101^. We assembled transcriptomes for each species using StringTie and identified hundreds of novel transcripts (with the prefix ‘MSTRG’), several of which were significantly upregulated after bacterial infection: 1-12 transcripts following *E. faecalis* infection and 0-16 following *P. rettgeri* infection depending on the species (Figure 6C and 6D, Table S7). We translated all valid ORFs on the sense strand of these transcripts and predicted signal peptides, identifying dozens of candidate secreted peptides (Figure 6E and 6F). These transcripts, mostly located in intergenic regions or introns and generally short in length, may encode novel AMPs and help resolve orthologous relationships in poorly annotated genomes. One example of these is the identified gene *Athelas-like* in *S. lebanonensis* described above (Figure 5A). Their strong induction upon infection, along with the challenge of detecting short, rapidly evolving sequences using BLAST or synteny search, underscore the value of de novo transcriptome assembly for AMP discovery and evolutionary analysis (Figure S5A-D).

To complement this differential expression-based approach, we developed a genome- and transcriptome-wide pipeline in *D. melanogaster* using machine learning (Figure S6 A, S6E). After benchmarking five prediction tools, we selected Macrel^102^ for its strong performance and applied it to genome- and transcript-derived ORFs, identifying 115 and 15 candidates with predicted signal peptides, respectively (Table S8). Many of these ORFs originated from non-genic or intronic regions, exhibited AMP-like physicochemical properties, and likely function independently of annotated host genes (Figure S6B-D, S6F). Together, these findings reveal a hidden repertoire of cryptic AMPs in the fly genome that are typically missed by conventional annotation and expression analysis.

### Transposable elements remain largely transcriptionally silent during bacterial infection

Given that transposable elements (TEs) may play a role in transcriptional regulations^103,104^, particularly under conditions of environmental stress or infection^105–107^, we investigated whether TEs contribute to gene regulation during immune responses to *E. faecalis* or *P. rettgeri*. In *D. melanogaster*, we reanalyzed RNA-seq data using a repeat-masked genome and a reference GTF file^108^ that includes annotated TEs. Of the 239 annotated transposable elements, only three showed significant differential expression following *E. faecalis* infection, and none were differentially expressed after *P. rettgeri* infection, compared to untreated groups (Table S9). Moreover, none of the differentially expressed TEs exhibited both high expression levels and substantial fold changes (average log_2_ fold-change = 1.09). These findings suggest that transposable elements are not broadly activated as part of the host transcriptional response to bacterial infection. Furthermore, the lack of widespread TE activation indicates that infection does not induce inappropriate TE expression, distinguishing the immune response from other stress conditions that may disrupt TE silencing. Overall, these results indicate that transposable elements remain largely transcriptionally silent during bacterial infection, indicating a limited role for TEs in regulating or responding to immune gene expression under these conditions.

## Discussion

In this study, we performed manual bacterial infection assays in five fly species spanning varying phylogenetic distances from *D. melanogaster*, using the Gram-positive pathogen *E. faecalis* and the Gram-negative pathogen *P. rettgeri*, both found in wild fruit flies. Quantitative RT-PCR of known AMP genes confirmed successful immune activation following infection. Survival monitoring over 30 days revealed distinct, species-specific infection outcomes. Although long-term survival was recorded, our analysis focused on the first two to three days post-infection - a critical period of acute immune activation and rapid pathogen proliferation. Bacterial load measurements indicated that variation in survival was primarily driven by differences in bacterial clearance rather than tolerance to comparable pathogen burdens. However, bacterial load was assessed at only a single time point (one day post-infection). Integrating longitudinal bacterial load measurements with transcriptomic profiling in future studies would provide a dynamic view of clearance mechanisms and their molecular regulation.

To understand the molecular basis of these divergent outcomes, we performed comparative transcriptomic profiling one day post-infection across all five species, including bacteria-infected, sterile-wounded, and untreated groups. While OrthoMCL effectively captured most orthologous relationships, incomplete annotations and high sequence divergence - especially among short peptides - prevented full recovery of all immune genes. As multi-copy families are difficult to compare across species, we restricted our analysis to single-copy orthologs, yielding a conservative but robust dataset representing core immune genes.

Using an expression divergence score (EDS) to quantify regulatory differences across species, we observed marked interspecies variations, reflecting diversification in host defense strategies and providing a foundation for understanding the genetic and regulatory bases of immune system evolution.

Previous work has emphasized lineage-specific immune genes essential to host defense. For example, *Drosomycin*, a Toll pathway-activated AMP in *D. melanogaster*, forms a seven-member gene family that arose through recent duplications and expanded independently in the *D. melanogaster* and *D. ananassae* subgroups^109,110^. Consistently, our analysis revealed lineage-specific genes that were transcriptionally induced upon infection, suggesting that each species employs distinct sets of immune effectors. Even conserved genes showed divergent expression patterns, highlighting regulatory evolution as a major force shaping immune diversity.

In *D. melanogaster*, both the number and magnitude of upregulated genes differed between *E. faecalis* and *P. rettgeri* infections. While technical variation may contribute, these patterns likely reflect biological differences: *E. faecalis* primarily activates the Toll pathway, which mediates defence against Gram-positive bacteria and fungi, whereas *P. rettgeri* is sensed largely through the Imd pathway, so the two infections engage partly non-overlapping regulons. Across species, infection predominantly triggered transcriptional activation rather than repression, supporting our focus on genes with elevated expression, especially AMP-like peptides.

Given the incomplete genome annotations of non-*melanogaster* species, total counts of differentially expressed genes should be interpreted cautiously. Some infection-induced genes may remain unannotated, and many lack functional characterization beyond *D. melanogaster* orthology. These limitations underscore the need for improved genome resources to achieve a fuller understanding of immune evolution. To partially address these gaps, we performed de novo transcript assembly and identified previously unannotated open reading frames encoding small, secreted peptides resembling AMPs.

Functional assays confirmed antimicrobial activity for several rapidly evolving genes, including *Daisho2*, *Athelas*, *Mtkl*, and *Athelas-like*, with minimum inhibitory concentrations (MICs) as low as 32 ug/mL. In particular, we found that *Daisho2,* previously characterized primarily for its antifungal activity, also possesses broad antibacterial activity against multiple bacterial strains. For *Athelas*, *Mtkl*, and *Athelas-like,* our assays revealed more general antimicrobial activities that had not been previously reported or experimentally tested. Because our MIC assays covered only a limited panel of bacterial and fungal strains pathogenic to flies and humans, expanding this panel could further clarify the breadth and mechanisms of their antimicrobial activities. While in vitro assays reflect intrinsic antimicrobial potential, in vivo efficacy is likely influenced by additional factors such as post-translational modifications and cofactor interactions, and thus requires further investigation.

Together, these results illustrate how rapidly evolving genes contribute to antimicrobial defense. By comparing five fly species, this study provides insight into immune system diversification across the Drosophilidae. Including more species in future work will help illuminate the evolutionary trajectory of immune responses and reveal additional AMP families.

Finally, using machine learning-based AMP prediction tools, we identified novel D. melanogaster peptides with physicochemical features resembling known AMPs, underscoring insect genomes as rich resources of cryptic antimicrobial peptides. As computational methods, particularly large language models, continue to improve, they may enable more accurate large-scale discovery of hidden AMPs across diverse taxa. Beyond AMPs, larger immune-related genes that function within complex protein-protein interaction networks^111^ remain to be explored. Although more difficult to characterize experimentally, these genes are critical to immune defense and represent promising targets for future studies on the evolution and integration of diverse immune components.

## Methods

### Fly strain preparation

Fly strains used in this study were obtained from multiple sources and subsequently maintained in the laboratory: *D. melanogaster* Canton-S (from Gaby Maimon, The Rockefeller University), *D. simulans* w501 (from David Begun, UC Davis), *D. ananassae* (from Vanessa Ruta, The Rockefeller University), *D. virilis* 15010.1051.47 (from Vanessa Ruta, The Rockefeller University), *S. lebanonensis* (from Andrew G. Clark, Cornell University). Flies were reared at 25°C, 60% relative humidity, on standard fly food under a 12:12-hour light/dark cycle. Because female flies are slightly more susceptible to bacterial infection^112^, we exclusively used females in all experiments. Flies were transferred to fresh vials within 0-3 days post-eclosion, and infections were carried out on healthy 5-7-day-old females. All anesthesia and infection procedures were performed at the same time of day to minimize circadian effects on gene expression.

### Bacteria preparation

*E. faecalis* (from Sean Brady, The Rockefeller University) and *P. rettgeri* (from Brian Lazzaro, Cornell University) were retrieved from glycerol stock, streaked onto LB agar plates, and incubated overnight at 37°C, then stored at 4°C. Liquid cultures were initiated from a single colony and grown overnight for 14-16 hours in 2 mL LB broth at 37°C with shaking. Bacteria cells were harvested by centrifugation, resuspended in sterile PBS, and adjusted to OD_600_ = 1 for injection. To account for body size differences, a two-fold higher dose was administered to *D. virilis* and *S. lebanonensis*. Body mass of 100 individuals from these two species and *D. melanogaster*, was measured to determine the scaling factor.

### Infection experiment

A 0.1 mm minutien pin was mounted into a pipette tip, which was sealed over a gas burner to secure the pin. Anesthetized flies on a CO2 pad were injected in the sternopleural plate of the right thorax, carefully avoiding wing and leg attachment sites. Two control groups were included: one injected with sterile 1x PBS and one left unmanipulated. Following injection, flies were transferred to standard food vials for recovery. Each group consisted of 10 flies per sample, with three biological replicates per condition. After 24 hours, 10 flies from each replicate were pooled, flash-frozen in 1.5 mL microcentrifuge tubes, and stored at −80°C for subsequent RNA extraction.

### Survival assay

Following infection, flies were maintained in groups of ten per vial on standard food. Survival was recorded daily, and flies were periodically transferred to fresh vials to minimize mortality unrelated to infection. Survival curves were generated using the Kaplan-Meier method^113,114^, and LT50 (median lethal time) was defined as the time point (in days) at which 50% of individuals had died. Cross symbols (“+”) on the survival curves indicate censored data due to loss of flies or uncertainty in survival tracking.

### Bacterial load quantification

Infection experiments for bacterial load quantification were performed as described above. Following infection, individual flies were homogenized in 250uL of sterile PBS using sterile pestles. For serial dilutions, 100 uL of homogenate from each fly was transferred to the first row of a 96-well plate, and 10 uL was serially diluted 10-fold across subsequent rows. From each dilution, 10 uL was plated onto pre-gridded LB agar plates. Plates were air-dried at room temperature until the liquid was absorbed, then incubated at 37℃ for 7-10 hours. Visible colonies were manually counted to estimate bacterial load, calculated as colony forming units per milliliter (CFU/mL). Dead flies were excluded from analysis to avoid confounding effects of post-mortem bacterial growth.

To evaluate the relationship between survival and bacterial load across species, we applied a Bayesian phylogenetic generalized least squares regression in PyMC^115^. Regression parameters were estimated using Markov chain Monte Carlo (MCMC) sampling, and posterior distributions were summarized with ArviZ^116^.

### RNA extraction

Total RNA was extracted using a standard TRIzol protocol with DNase treatment, and concentration and purity were assessed using a NanoDrop spectrophotometer. The same RNA samples were used for both quantitative RT-PCR and RNA-seq library preparation (described below).

### Quantitative RT-PCR

Reverse transcription was performed using the PrimeScript RT Reagent Kit (TaKaRa, Cat#RR037A). Quantitative PCR was conducted with PowerUp SYBR Green Master Mix (ThermoFisher, Cat#A25741) on selected immune-related genes, including *Drosocin*, *Cecropin*, and *Diptericin*. *Ribosomal protein l32* (*Rpl32*) served as the reference gene for normalization^117,118^. Because of incomplete gene annotation or the absence of orthologs in some species, the set of AMP genes analyzed varied among species.

Fluorescence signals were detected on a QuantStudio 5 Real-Time PCR system, and fold changes in gene expression were calculated using the ΔCt method. All experiments have three or more biological replicates.

### RNA sequencing and data analysis

RNA-seq libraries were prepared using the NEBNext Ultra Ⅱ Directional RNA Library Prep Kit for Illumina (New England BioLabs, #E7760S/L, #E7765S/L) and sequenced by Novogene Corporation Inc (www.novogene.com). Libraries were run on the Illumina NovaSeq platform with 150 bp paired-end reads, generating an average of 37.9 million reads per library. Quality control of raw sequencing data was performed with FastQC^119^ and MultiQC^120^, and adapter trimming was conducted with trimmomatic^121^. Reference genomes were obtained from FlyBase (*D. melanogaster*: dmel-r6.48) and NCBI (for *D. simulans* - GCF_016746395.2_Prin_Dsim_3.1, *D. ananassae* - GCF_017639315.1_ASM1763931v2, *D. virilis* - GCF_003285735.1_DvirRS2, and *S. lebanonensis* - GCF_003285725.1_SlebRS2).

Reads were aligned to the genomes using Hisat2^122^ and assembled into transcriptomes with Stringtie^123^. Transcript and gene abundance was quantified with Stringtie, and raw counts were obtained using HTseq^124^. Differential expression analysis was performed in DESeq2^125^, with genes classified as significantly up- or downregulated at thresholds of |log_2_ fold-change| > 1 and adjusted *p* < 0.05. For visualization, volcano plots were generated using shrunken fold changes estimates from the *ashr* algorithm^126^ in R, which improves reliability by reducing exaggerated fold changes for genes with low counts or high variance.

### Gene conservation and expression divergence

Phylogenetic relationships across the five species were taken from previous studies^127,128^ for phylogenetic tree construction and downstream analysis. Orthologous gene groups were identified using OrthoMCL, which conducts an all-vs-all BLASTP search across high-quality protein sequences from the five species.

Protein-level orthologs were then mapped to gene ortholog groups based on genome annotations. From this analysis, we identified 10,440 conserved genes present in all five species, regardless of copy number, of which 9,296 were single-copy in each species.

To quantify interspecific variation in gene regulation, we calculated an expression divergence score (EDS) for each ortholog group from log_2_ fold-change values following infection relative to untreated controls:

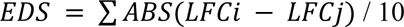

where *LFCi* and *LFCj* are the log₂ fold-change values in two different species, across all ten pairwise species comparisons. The final EDS was obtained as the average of these absolute differences. A higher EDS indicates greater divergence in infection-induced gene expression across species.

Pagel’s lambda quantifies the extent to which trait variance is explained by phylogeny, providing a measure of the phylogenetic signal in trait evolution. We estimated lambda using the *ape* and *phytools* packages in R, based on log_2_ fold-change in gene expression between untreated and bacteria-infected groups. Only genes with TPM > 1 across all five species post-infection were included. Genes were classified as upregulated (significantly upregulated in at least one species) or other (not significantly upregulated in any species), and those with comparable expression levels were selected for analysis using simulated annealing^129^.

To assess expression conservation, we calculated an induction ratio, defined as the proportion of genes upregulated in more than half of the species relative to those upregulated in at least one species. A higher induction ratio indicates greater conservation of transcriptional responses to infection.

### Mantel test

Immune-response similarity between species was quantified using the Jaccard distance^130^, calculated from binary profiles indicating whether a gene was significantly upregulated in each species (1 = upregulated, 0 = not upregulated). The correlation between the immune-response similarity matrix and the phylogenetic distance matrix was evaluated using the Mantel test^131^ with Pearson’s correlation and 999 permutations to assess significance.

### Analysis of upregulated genes in the resistant group

To identify genes specifically upregulated in the resistant group, we first applied a log2 fold-change threshold to one-to-one orthologs: genes with log_2_ fold-change > 2 in resistant species and < 2 in susceptible species, based on comparisons between untreated and bacteria-infected conditions. In parallel, a count matrix restricted to bacteria-infected samples was analyzed with DESeq2 to identify genes highly expressed in the resistant group (baseMean > 500, significantly upregulated relative to the susceptible group). Results from both approaches were merged and filtered to retain only proteins shorter than 200 amino acids and to remove redundant entries.

### Identification of rapidly evolving, infection-induced small genes and tissue-specific expression

Building on the RNA-seq differential expression results, we identified small, rapidly evolving genes that were strongly induced by bacterial infection. Rapidly evolving genes were defined as conserved genes upregulated in a subset - but not all - species, and non-conserved genes upregulated following bacterial infection. High expression in immune-related tissues was determined based on gene expression levels in the fat body and/or hemocytes, using publicly available datasets from the Fly Cell Atlas single-cell RNA-seq^132^ and FlyAtlas2 Anatomy RNA-seq^93^ (female) databases.

### Unannotated open reading frame identification

To identify putative open reading frames in unannotated transcripts, transcriptomes were assembled using Stringtie. We used NCBI ORFfinder^133^ to detect all possible ORFs on the correct strand of each transcript. To ensure novelty, we used genomic coordinates to confirm that none of the identified ORFs overlapped with annotated exons. We then used SignalP6^134^ to predict N-terminal signal peptides, indicative of potential secretion.

### Evolutionary analysis of AMPs

To identify AMP genes across species, we performed TBLASTN^135^ to search against the reference genomes of each species using the protein sequences from *D. melanogaster* or preidentified orthologs. For genes that were difficult to recover using TBLASTN, we identified the orthologous region using more conserved flanking genes, which were identified using a similar strategy. After extracting the sequence of orthologous regions, the potential gene coding sequences were identified by GeneWise^136^.

To determine the phylogenetic relationship of *Athelas* homologs, we aligned the deduced amino acid sequences of mature regions using MAFFT algorithm E-INS-i^137^ and generated nucleotide sequence alignment based on the protein sequence alignment using PAL2NAL^138^. The best-fit model HKY+G was selected by MrModelTest 2.4^139^. Phylogenetic inference was implemented using MrBayes 3.2.7^140^. *Mtkl* from *D. melanogaster* and *P. okadai* were used as outgroups.

### Peptide synthesis

Candidate peptides were synthesized at ≥ 95% purity by LifeTein, LLC (www.lifetein.com) using solid-phase peptide synthesis and verified by mass spectrometry. Peptide sequences excluded the signal peptide region of the parent gene, and selected peptides were chemically modified by N-terminal acetylation and C-terminal amidation to enhance stability. Lyophilized peptides were dissolved in water, quantified, and stored at −20°C until use.

### Antimicrobial activity assay

In addition to *E. faecalis* and *P. rettgeri* described above, the following bacterial and fungal strains were used in the assay: *Staphylococcus aureus*, *Acinetobacter baumannii*, *Enterobacter cloacae*, *Pseudomonas aeruginosa*, *Klebsiella pneumoniae*, *Enterococcus faecium*, *Bacillus subtilis*, and *Candida albicans* (from Sean Brady, The Rockefeller University). Liquid bacterial cultures were adjusted to an optical density (OD_600_) of 1, diluted to 0.1, and further diluted 100-fold to approximately 0.001. Peptide solutions were serially diluted two-fold from the highest to lowest concentration. Equal volumes of bacterial suspension and peptide solution were combined in designated wells of a 96-well plate, with the top and bottom rows left empty to minimize edge effects. Plates were incubated statically at 37°C or 30°C for 20-24 hours.

After incubation, wells were examined with a microscope, and resazurin - a redox-sensitive dye - was added to assess bacterial viability. Viable cells reduce resazurin, producing a color change from blue to pink, which was visually assessed and recorded after 2 hours.

### AMP prediction using machine learning

We used the same genome references and transcriptomes as in the RNA-seq analysis. To identify potential AMPs, we implemented a multi-step computational pipeline (Figure S6A). First, we benchmarked five models from three AMP prediction tools CAMPR4^141^, Antimicrobial Peptide Scanner vr.2^142^, and Macrel^102^ using a curated dataset of AMPs and non-AMPs compiled from public AMP databases APD3^143^ and LAMP2^144^, and a general protein database Uniprot^145^ (Table S8). Macrel showed the best performance, with an accuracy of 0.90, precision of 0.98, specificity of 0.99, and recall of 0.66, and was selected for subsequent predictions.

We applied NCBI ORFfinder to detect all possible ORFs from both genomes and transcriptomes. ORFs were filtered for uniqueness and length (< 200 amino acids). Candidate AMPs were predicted using Macrel (probability cutoff > 0.5). To exclude annotated proteins, we performed BLASTP against known protein annotations, using thresholds of bit score > 100 and query match percentage > 40%.

Signal peptides were predicted using SignalP6, and genomic coordinates of candidate ORFs were cross-referenced with gene annotation files using a custom Python script to identify overlaps with known genomic features. Finally, candidate ORFs were categorized based on their location relative to annotated genes (e.g. intergenic, intronic, or overlapping).

## Code and data availability

RNA-sequencing data is submitted to NCBI BioProject under the accession number PRJNA1391518. Custom Python and R scripts used for data analysis in this study, along with the corresponding datasets, are available on the project’s Github repository: https://github.com/LiZhaoLab/Evolution_immune.

## Acknowledgements

We thank members of the Zhao laboratory for helpful discussions. We thank Dr. Sean Brady of The Rockefeller University and Dr. Brian Lazzaro of Cornell University for providing the bacterial strains. We also thank the Rockefeller University High Performance Computing Center and the Genome Resource Center for computational and genomic support, respectively.

## Supplementary information

**Figure S1.**
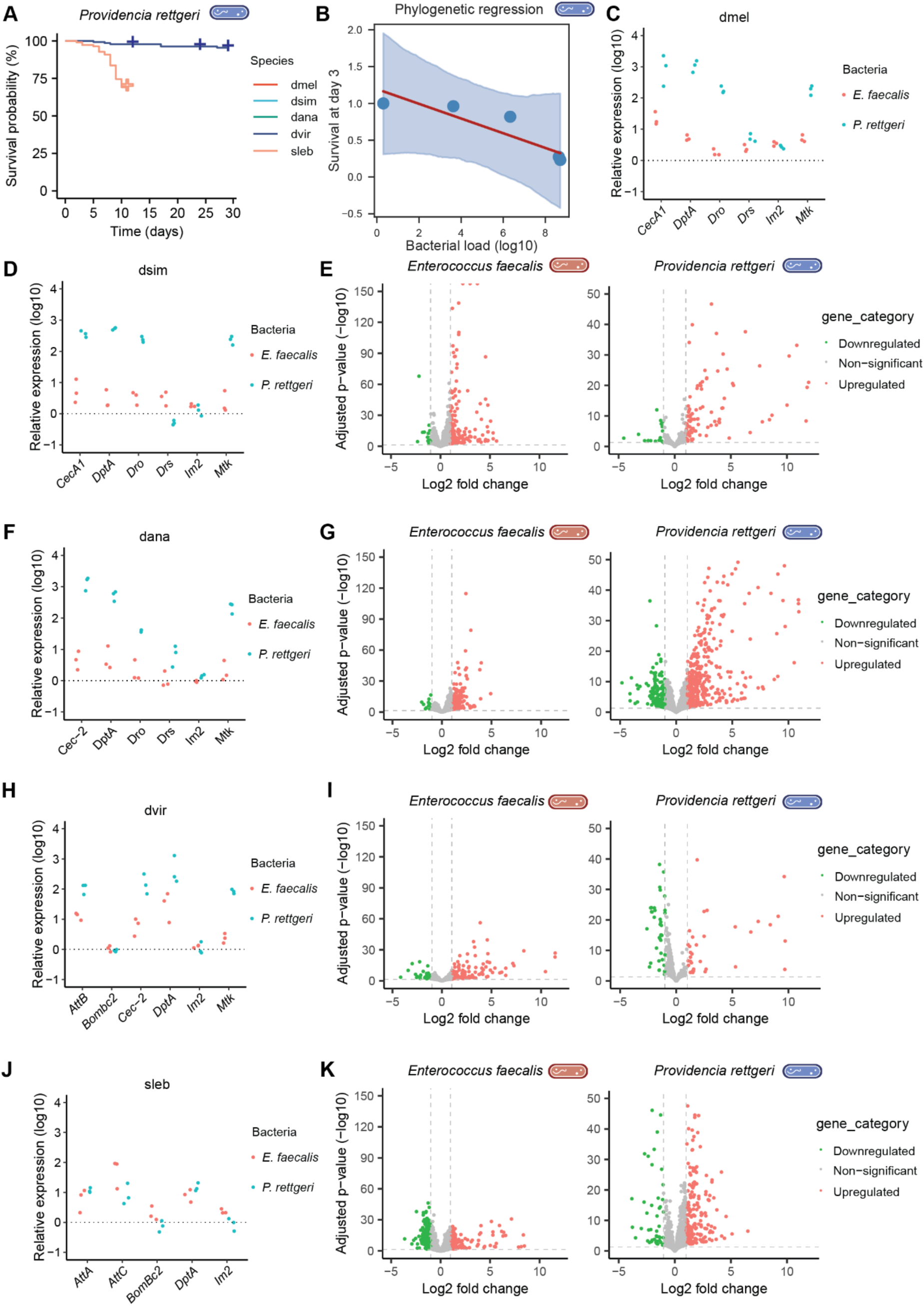
Survival was maintained under control conditions, and immune responses were induced in all five species. A) Body size-normalized survival curves confirmed that the observed variance in mortality is due to bacterial infection. B) Scatter plot showing the relationship between log₁₀-transformed bacterial load at day 1 and survival at day 3 following *P. rettgeri* infection. Each point represents the median bacterial load and mean survival for a species. A Bayesian phylogenetic generalized least squares regression estimated a posterior mean correlation coefficient of ρ = - 0.875, indicating that species with higher bacterial loads tend to exhibit lower survival. The shaded region represents the 95% CI of the regression fit. C, D, F, H, J) Quantitative RT-PCR of known antimicrobial peptide (AMP) genes showed strong upregulation after bacterial infection compared to sterile-wounded controls. E, G, I, K) The presence of both upregulated and downregulated genes further confirmed that immune responses were robustly induced in all five species.

**Figure S2.**
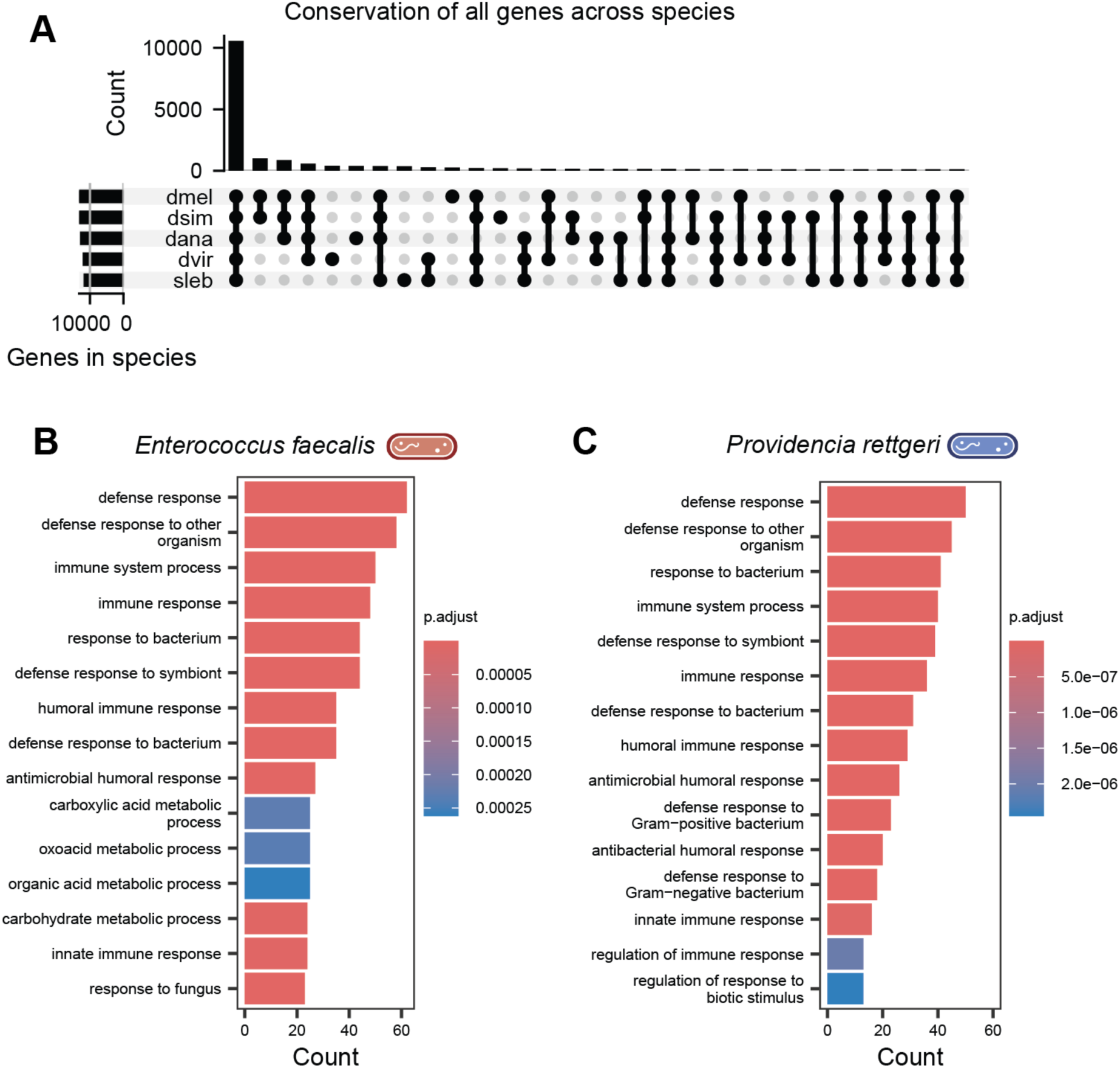
Orthologous gene groups and species-specific immune expression profiles. A) OrthoMCL analysis identified 10,440 orthologous gene groups shared across the five species. B-C) Gene Ontology enrichment analysis of upregulated gene in *D. melanogaster* revealed top enriched terms associated with immune-related functions.

**Figure S3.**
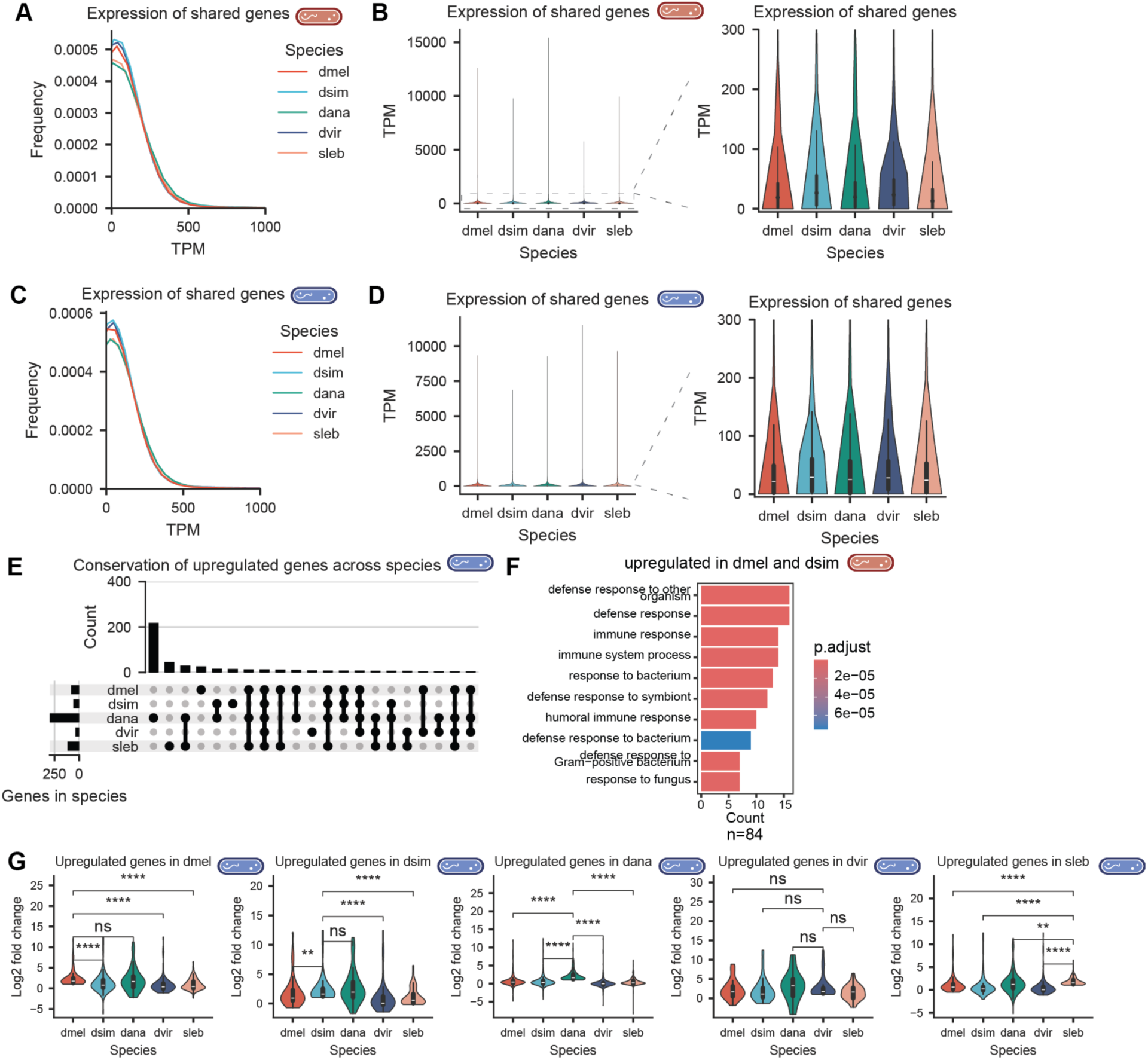
Conserved genes exhibit similar expression patterns in general, but regulatory divergence follows *P. rettgeri* infection. A-B) Expression patterns of single-copy conserved orthologs following *E. faecalis* infections across species. Most genes show broadly similar expression profiles across species. Distributions of TPM values for conserved genes are shown for each species, with a zoomed-in view for genes with TPM < 300 to highlight lower-expression genes. C-D) One-to-one conserved orthologs generally showed similar expression profiles across species after *P. rettger* infection. E) However, upregulation of gene expression was not conserved, indicating divergence in infection-induced regulation. F) Genes co-upregulated in *D. melanogaster* and in *D. simulans* were enriched for immune-related GO terms. G) Genes upregulated in a given focal species exhibited significantly higher fold changes in that species compared to their orthologs in other species, with a few exceptions (ns = not significant; *, *p* < 0.05; **, *p* < 0.01; ***, *p* < 0.001; ****, *p* < 0.0001; Mann–Whitney U test with FDR-BH correction).

**Figure S4.**
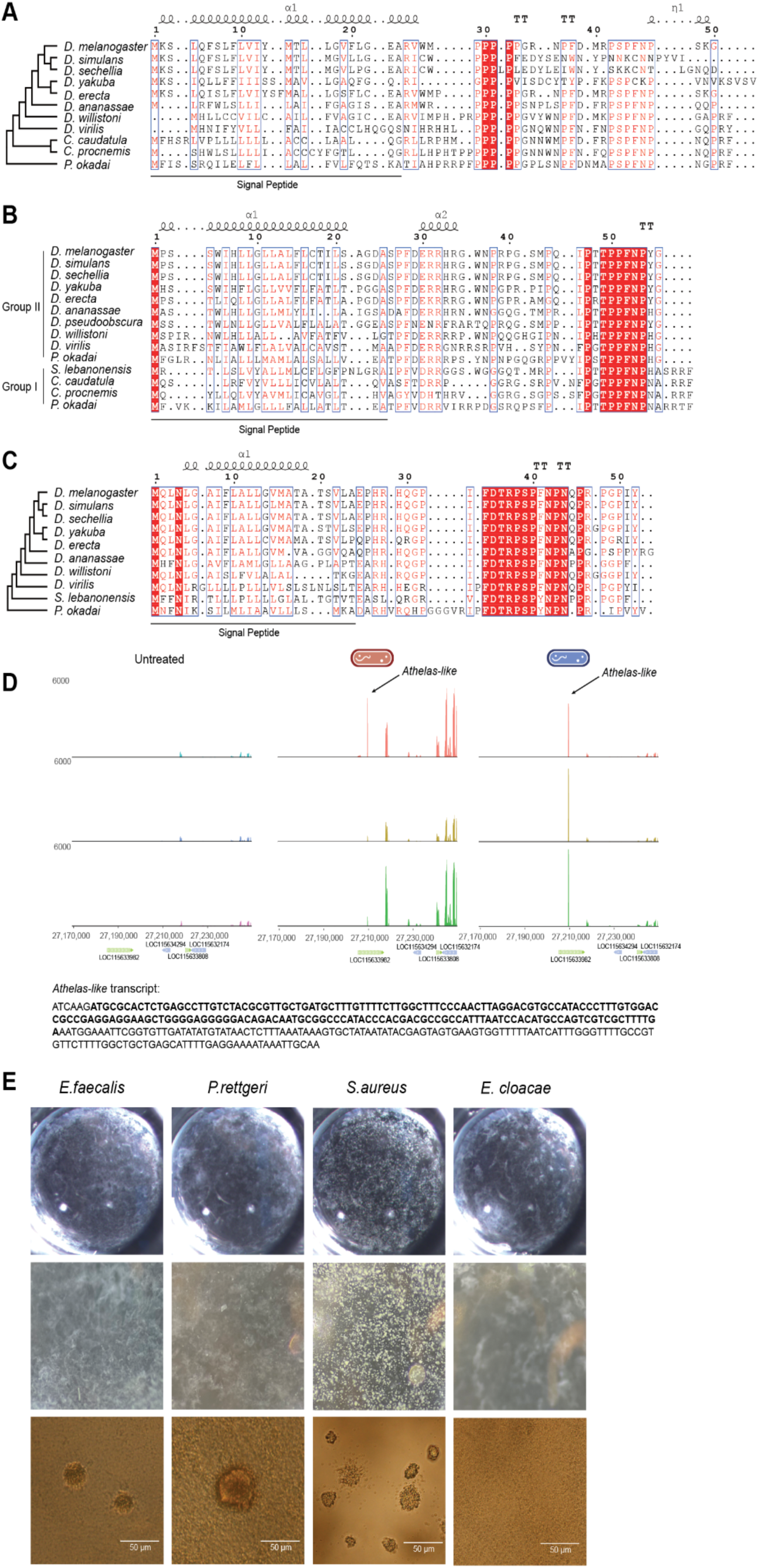
Alignments and representative phenotypes of bacteria treated with Daisho2. Protein sequence alignments of Mtkl (A), Athelas and Athelas-like (B), and Mtk (C). D) RNA-seq coverage of *Athelas-like* in the untreated group (left), after *E. faecalis* infection (middle), and after *P. rettgeri* infection (right).The transcript sequence of *Athelas-like* is shown (bottom), with the coding sequence highlighted in bold. E) Representative images of bacterial cultures of *E. faecalis*, *P. rettgeri*, *S. aureus*, and *E. cloacae* incubated with the peptide Daisho2, showing characteristic growth phenotypes following treatment.

**Figure S5.**
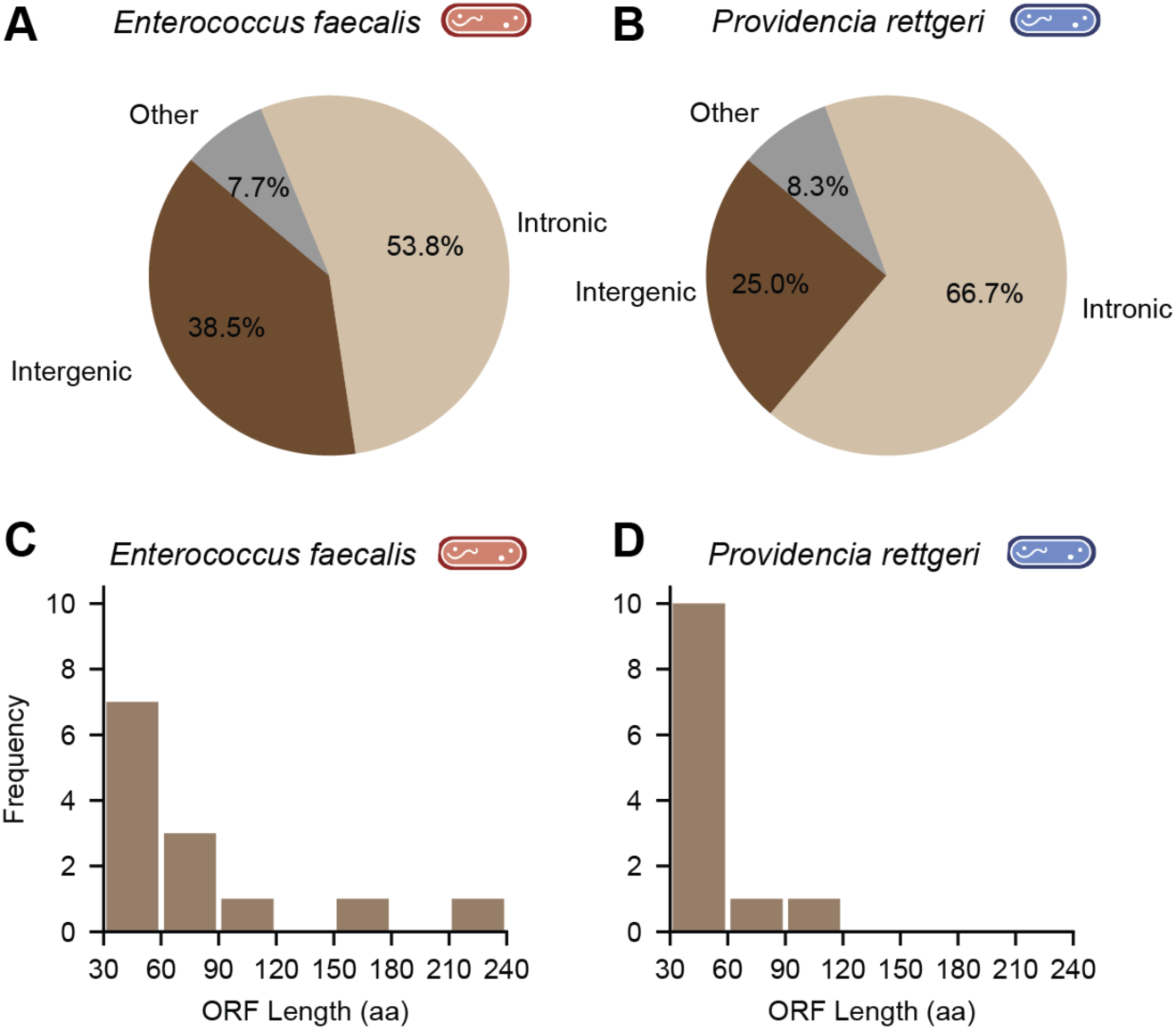
Genomic origins and length distributions of ORFs with predicted signal peptides. A-B) Most ORFs containing signal peptides are located in intergenic or intronic regions, rather than annotated exons. C-D) These ORFs are generally short, consistent with typical characteristics of antimicrobial peptides.

**Figure S6.**
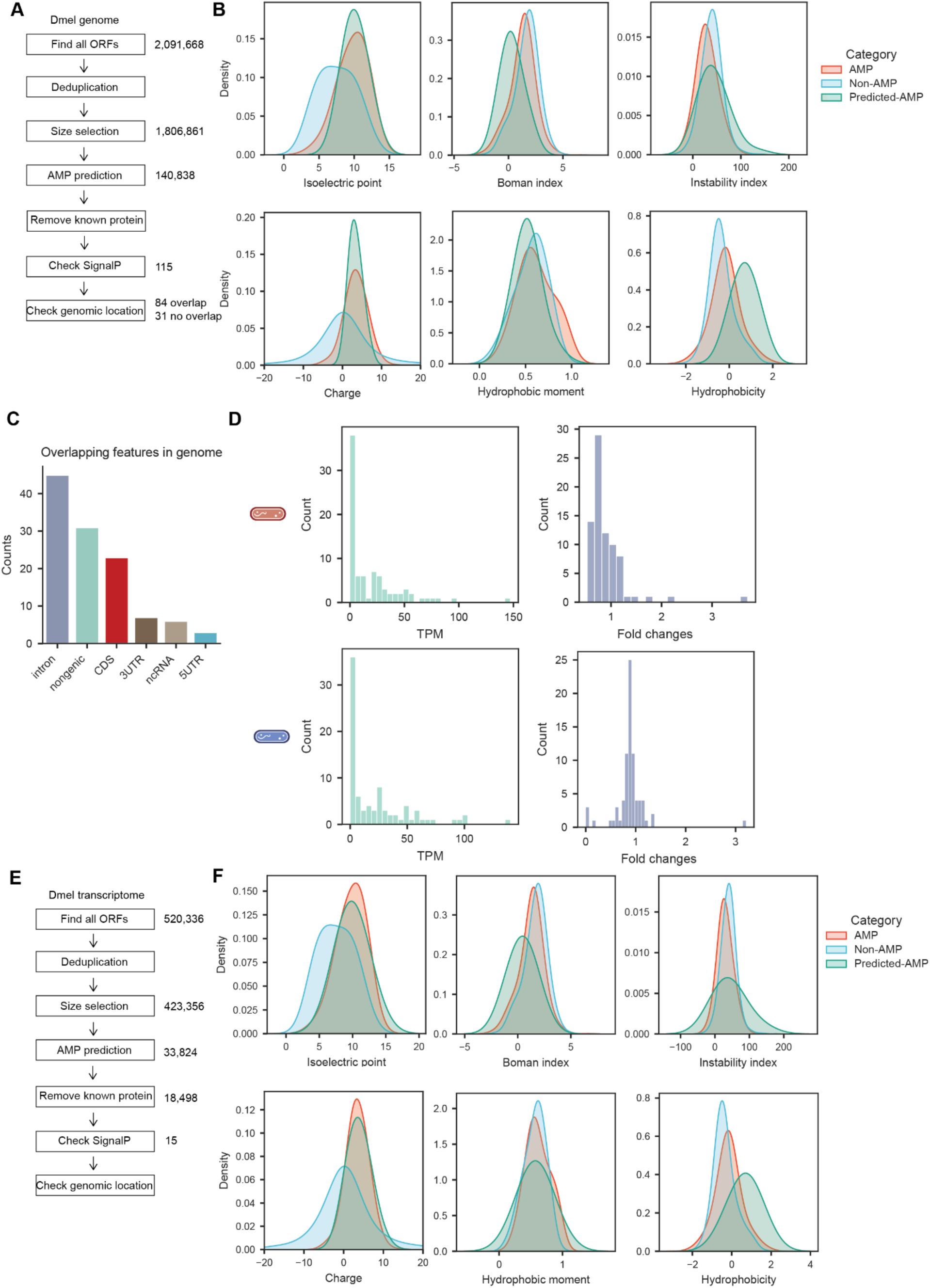
Putative antimicrobial peptides (AMPs) identified using machine learning-based prediction tools. A) Schematic overview of the computational pipeline used to identify AMP-like peptides from the *D. melanogaster* genome. B) Candidate peptides exhibit physicochemical properties similar to known AMPs and distinct from non-AMP proteins. C) Among the 115 genome-derived AMP candidates with predicted signal peptides, most are located in intronic or intergenic regions. D) Host genes overlapping with candidate ORFs tend to be lowly expressed and downregulated. E) Schematic of the pipeline used to identify AMP-like peptides from the *D. melanogaster* transcriptome. F) Physicochemical profiles of transcriptome-derived peptides resemble those of known AMPs and are distinct from non-AMP proteins.

## References

1. Tzou, P. How Drosophila combats microbial infection: a model to study innate immunity and host– pathogen interactions. Curr. Opin. Microbiol. 5, 102–110 (2002).

2. Hultmark, D. Drosophila immunity: paths and patterns. Curr. Opin. Immunol. 15, 12–19 (2003).

3. Hoffmann, J. A. The immune response of Drosophila. Nature 426, 33–38 (2003).

4. Brennan, C. A. & Anderson, K. V. *DROSOPHILA*: The Genetics of Innate Immune Recognition and Response. Annu. Rev. Immunol. 22, 457–483 (2004).

5. De Gregorio, E. The Toll and Imd pathways are the major regulators of the immune response in Drosophila. EMBO J. 21, 2568–2579 (2002).

6. Lemaitre, B. & Hoffmann, J. The Host Defense of *Drosophila melanogaster*. Annu. Rev. Immunol. 25, 697–743 (2007).

7. Imler, J.-L. & Bulet, P. Antimicrobial Peptides in Drosophila: Structures,Activities and Gene Regulation. in Chemical Immunology and Allergy (eds Kabelitz, D. & Schröder, J.-M.) vol. 86 1–21 (S. Karger AG, 2005).

8. Mookherjee, N., Anderson, M. A., Haagsman, H. P. & Davidson, D. J. Antimicrobial host defence peptides: functions and clinical potential. Nat. Rev. Drug Discov. 19, 311–332 (2020).

9. Lazzaro, B. P., Zasloff, M. & Rolff, J. Antimicrobial peptides: Application informed by evolution. Science 368, eaau5480 (2020).

10. Lin, S. J. H., Cohen, L. B. & Wasserman, S. A. Effector specificity and function in Drosophila innate immunity: Getting AMPed and dropping Boms. PLOS Pathog. 16, e1008480 (2020).

11. Hanson, M. A. & Lemaitre, B. New insights on Drosophila antimicrobial peptide function in host defense and beyond. Curr. Opin. Immunol. 62, 22–30 (2020).

12. Schlenke, T. A. & Begun, D. J. Natural Selection Drives Drosophila Immune System Evolution. Genetics 164, 1471–1480 (2003).

13. Keebaugh, E. S. & Schlenke, T. A. Insights from natural host–parasite interactions: The Drosophila model. Dev. Comp. Immunol. 42, 111–123 (2014).

14. Duxbury, E. M. et al. Host-pathogen coevolution increases genetic variation in susceptibility to infection. eLife 8, e46440 (2019).

15. Hill, T., Koseva, B. S. & Unckless, R. L. The Genome of Drosophila innubila Reveals Lineage-Specific Patterns of Selection in Immune Genes. Mol. Biol. Evol. 36, 1405–1417 (2019).

16. Dhakad, P. & Obbard, D. J. Predictors of protein evolution in the drosophilid immune system. Genome Biol. Evol. evag034 (2026) doi:10.1093/gbe/evag034.

17. Obbard, D. J., Welch, J. J., Kim, K.-W. & Jiggins, F. M. Quantifying Adaptive Evolution in the Drosophila Immune System. PLoS Genet. 5, e1000698 (2009).

18. Jiggins, F. M. & Kim, K. W. A screen for immunity genes evolving under positive selection in *Drosophila*. J. Evol. Biol. 20, 965–970 (2007).

19. Sackton, T. B. et al. Dynamic evolution of the innate immune system in Drosophila. Nat. Genet. 39, 1461–1468 (2007).

20. Unckless, R. L., Howick, V. M. & Lazzaro, B. P. Convergent Balancing Selection on an Antimicrobial Peptide in Drosophila. Curr. Biol. 26, 257–262 (2016).

21. Chapman, J. R., Hill, T. & Unckless, R. L. Balancing Selection Drives the Maintenance of Genetic Variation in Drosophila Antimicrobial Peptides. Genome Biol. Evol. 11, 2691–2701 (2019).

22. Hanson, M. A., Grollmus, L. & Lemaitre, B. Ecology-relevant bacteria drive the evolution of host antimicrobial peptides in *Drosophila*. Science 381, eadg5725 (2023).

23. Hanson, M. A. et al. Synergy and remarkable specificity of antimicrobial peptides in vivo using a systematic knockout approach. eLife 8, e44341 (2019).

24. Vinkler, M. et al. Understanding the evolution of immune genes in jawed vertebrates. J. Evol. Biol. 36, 847–873 (2023).

25. Long, M., Betrán, E., Thornton, K. & Wang, W. The origin of new genes: glimpses from the young and old. Nat. Rev. Genet. 4, 865–875 (2003).

26. Levine, M. T., Jones, C. D., Kern, A. D., Lindfors, H. A. & Begun, D. J. Novel genes derived from noncoding DNA in *Drosophila melanogaster* are frequently X-linked and exhibit testis-biased expression. Proc. Natl. Acad. Sci. 103, 9935–9939 (2006).

27. Chen, S., Krinsky, B. H. & Long, M. New genes as drivers of phenotypic evolution. Nat. Rev. Genet. 14, 645–660 (2013).

28. Zhao, L., Svetec, N. & Begun, D. J. De Novo Genes. Annu. Rev. Genet. 58, 211–232 (2024).

29. Zhuang, X., Yang, C., Murphy, K. R. & Cheng, C.-H. C. Molecular mechanism and history of non-sense to sense evolution of antifreeze glycoprotein gene in northern gadids. Proc. Natl. Acad. Sci. 116, 4400–4405 (2019).

30. Peng, J. & Zhao, L. The origin and structural evolution of de novo genes in Drosophila. Nat. Commun. 15, 810 (2024).

31. Torres, M. D. T. et al. Mining for encrypted peptide antibiotics in the human proteome. Nat. Biomed. Eng. 6, 67–75 (2021).

32. Cesaro, A. et al. Synthetic Antibiotic Derived from Sequences Encrypted in a Protein from Human Plasma. ACS Nano 16, 1880–1895 (2022).

33. Wan, F., Torres, M. D. T., Peng, J. & De La Fuente-Nunez, C. Deep-learning-enabled antibiotic discovery through molecular de-extinction. Nat. Biomed. Eng. 8, 854–871 (2024).

34. Dhakad, P., Newman, D. & Obbard, D. J. Transcriptomic analysis of non-model Drosophilidae reveals novel AMP candidates. BMC Biol. 24, 62 (2026).

35. Jackson, T. J. et al. Isolation of insect pathogenic bacteria, *Providencia rettgeri*, from *Heterorhabditis* spp. J. Appl. Bacteriol. 78, 237–244 (1995).

36. Franz, C. M. A. P. et al. Incidence of Virulence Factors and Antibiotic Resistance among Enterococci Isolated from Food. Appl. Environ. Microbiol. 67, 4385–4389 (2001).

37. Yoh, M. et al. Importance of Providencia species as a major cause of travellers’ diarrhoea. J. Med. Microbiol. 54, 1077–1082 (2005).

38. Gardini, F. et al. Modeling the Aminogenic Potential of *Enterococcus faecalis* EF37 in Dry Fermented Sausages through Chemical and Molecular Approaches. Appl. Environ. Microbiol. 74, 2740–2750 (2008).

39. Hanchi, H., Mottawea, W., Sebei, K. & Hammami, R. The Genus Enterococcus: Between Probiotic Potential and Safety Concerns—An Update. Front. Microbiol. 9, 1791 (2018).

40. Cox, C. R. & Gilmore, M. S. Native Microbial Colonization of *Drosophila melanogaster* and Its Use as a Model of *Enterococcus faecalis* Pathogenesis. Infect. Immun. 75, 1565–1576 (2007).

41. Troha, K., Im, J. H., Revah, J., Lazzaro, B. P. & Buchon, N. Comparative transcriptomics reveals CrebA as a novel regulator of infection tolerance in D. melanogaster. PLOS Pathog. 14, e1006847 (2018).

42. Kylsten, P., Samakovlis, C. & Hultmark, D. The cecropin locus in Drosophila; a compact gene cluster involved in the response to infection. EMBO J. 9, 217–224 (1990).

43. Wicker, C. et al. Insect immunity. Characterization of a Drosophila cDNA encoding a novel member of the diptericin family of immune peptides. J. Biol. Chem. 265, 22493–22498 (1990).

44. Lemaitre, B. et al. A recessive mutation, immune deficiency (imd), defines two distinct control pathways in the Drosophila host defense. Proc. Natl. Acad. Sci. 92, 9465–9469 (1995).

45. Lemaitre, B., Nicolas, E., Michaut, L., Reichhart, J.-M. & Hoffmann, J. A. The Dorsoventral Regulatory Gene Cassette spätzle/Toll/cactus Controls the Potent Antifungal Response in Drosophila Adults. Cell 86, 973–983 (1996).

46. Galac, M. R. & Lazzaro, B. P. Comparative pathology of bacteria in the genus Providencia to a natural host, Drosophila melanogaster. Microbes Infect. 13, 673–683 (2011).

47. Lazzaro, B. P., Sackton, T. B. & Clark, A. G. Genetic Variation in *Drosophila melanogaster* Resistance to Infection: A Comparison Across Bacteria. Genetics 174, 1539–1554 (2006).

48. Louie, A., Song, K. H., Hotson, A., Thomas Tate, A. & Schneider, D. S. How Many Parameters Does It Take to Describe Disease Tolerance? PLOS Biol. 14, e1002435 (2016).

49. Duneau, D. et al. Stochastic variation in the initial phase of bacterial infection predicts the probability of survival in D. melanogaster. eLife 6, e28298 (2017).

50. Chambers, M. C., Jacobson, E., Khalil, S. & Lazzaro, B. P. Consequences of chronic bacterial infection in Drosophila melanogaster. PLOS ONE 14, e0224440 (2019).

51. Fuse, N. et al. Transcriptome features of innate immune memory in Drosophila. PLOS Genet. 18, e1010005 (2022).

52. Cabrera, K., Hoard, D. S., Gibson, O., Martinez, D. I. & Wunderlich, Z. Drosophila immune priming to Enterococcus faecalis relies on immune tolerance rather than resistance. PLOS Pathog. 19, e1011567 (2023).

53. Ekengren, S. & Hultmark, D. A Family of Turandot-Related Genes in the Humoral Stress Response of Drosophila. Biochem. Biophys. Res. Commun. 284, 998–1003 (2001).

54. Clemmons, A. W., Lindsay, S. A. & Wasserman, S. A. An Effector Peptide Family Required for Drosophila Toll-Mediated Immunity. PLOS Pathog. 11, e1004876 (2015).

55. Amstrup, A. B., Bæk, I., Loeschcke, V. & Givskov Sørensen, J. A functional study of the role of Turandot genes in Drosophila melanogaster: An emerging candidate mechanism for inducible heat tolerance. J. Insect Physiol. 143, 104456 (2022).

56. Xu, R. et al. The Toll pathway mediates *Drosophila* resilience to *Aspergillus* mycotoxins through specific Bomanins. EMBO Rep. 24, e56036 (2023).

57. Lindsay, S. A., Lin, S. J. H. & Wasserman, S. A. Short-Form Bomanins Mediate Humoral Immunity in **Drosophila**. J. Innate Immun. 10, 306–314 (2018).

58. Galko, M. J. & Krasnow, M. A. Cellular and Genetic Analysis of Wound Healing in Drosophila Larvae. PLoS Biol. 2, e239 (2004).

59. Razzell, W., Wood, W. & Martin, P. Swatting flies: modelling wound healing and inflammation in *Drosophila*. Dis. Model. Mech. 4, 569–574 (2011).

60. Lin, S. J. H., Fulzele, A., Cohen, L. B., Bennett, E. J. & Wasserman, S. A. Bombardier Enables Delivery of Short-Form Bomanins in the Drosophila Toll Response. Front. Immunol. 10, 3040 (2020).

61. Liu, S. et al. A tissue injury sensing and repair pathway distinct from host pathogen defense. Cell 186, 2127–2143.e22 (2023).

62. Smith, B. R., Patch, K. B., Gupta, A., Knoles, E. M. & Unckless, R. L. The genetic basis of variation in immune defense against Lysinibacillus fusiformis infection in Drosophila melanogaster. PLOS Pathog. 19, e1010934 (2023).

63. Steiner, H. Peptidoglycan recognition proteins: on and off switches for innate immunity. Immunol. Rev. 198, 83–96 (2004).

64. Shaka, M., Arias-Rojas, A., Hrdina, A., Frahm, D. & Iatsenko, I. Lipopolysaccharide-mediated resistance to host antimicrobial peptides and hemocyte-derived reactive-oxygen species are the major Providencia alcalifaciens virulence factors in Drosophila melanogaster. PLOS Pathog. 18, e1010825 (2022).

65. Li, L., Stoeckert, C. J. & Roos, D. S. OrthoMCL: Identification of Ortholog Groups for Eukaryotic Genomes. Genome Res. 13, 2178–2189 (2003).

66. Dimarcq, J. et al. Characterization and transcriptional profiles of a *Drosophila* gene encoding an insect defensin: A study in insect immunity. Eur. J. Biochem. 221, 201–209 (1994).

67. Tzou, P., Reichhart, J.-M. & Lemaitre, B. Constitutive expression of a single antimicrobial peptide can restore wild-type resistance to infection in immunodeficient *Drosophila* mutants. Proc. Natl. Acad. Sci. 99, 2152–2157 (2002).

68. Koehbach, J. Structure-Activity Relationships of Insect Defensins. Front. Chem. 5, 45 (2017).

69. Dionne, M. S., Pham, L. N., Shirasu-Hiza, M. & Schneider, D. S. Akt and foxo Dysregulation Contribute to Infection-Induced Wasting in Drosophila. Curr. Biol. 16, 1977–1985 (2006).

70. Darby, A. M. & Lazzaro, B. P. Interactions between innate immunity and insulin signaling affect resistance to infection in insects. Front. Immunol. 14, 1276357 (2023).

71. Lynch, M. METHODS FOR THE ANALYSIS OF COMPARATIVE DATA IN EVOLUTIONARY BIOLOGY. Evolution 45, 1065–1080 (1991).

72. Pagel, M. D. A method for the analysis of comparative data. J. Theor. Biol. 156, 431–442 (1992).

73. Freckleton, R. P., Harvey, P. H. & Pagel, M. Phylogenetic Analysis and Comparative Data: A Test and Review of Evidence. Am. Nat. 160, 712–726 (2002).

74. Sun, H. et al. Varying phylogenetic signal in susceptibility to four bacterial pathogens across species of Drosophilidae. Proc. R. Soc. B Biol. Sci. 292, 20242239 (2025).

75. Paparazzo, F., Tellier, A., Stephan, W. & Hutter, S. Survival Rate and Transcriptional Response upon Infection with the Generalist Parasite Beauveria bassiana in a World-Wide Sample of Drosophila melanogaster. PLOS ONE 10, e0132129 (2015).

76. Zmora, N., Bashiardes, S., Levy, M. & Elinav, E. The Role of the Immune System in Metabolic Health and Disease. Cell Metab. 25, 506–521 (2017).

77. Kedia-Mehta, N. & Finlay, D. K. Competition for nutrients and its role in controlling immune responses. Nat. Commun. 10, 2123 (2019).

78. Troha, K. & Ayres, J. S. Metabolic Adaptations to Infections at the Organismal Level. Trends Immunol. 41, 113–125 (2020).

79. Deshpande, R., Lee, B. & Grewal, S. S. Enteric bacterial infection in *Drosophila* induces whole-body alterations in metabolic gene expression independently of the immune deficiency signaling pathway. G3 GenesGenomesGenetics 12, jkac163 (2022).

80. Darby, A. M., Keith, S. A., Kalukin, A. A. & Lazzaro, B. P. Chronic bacterial infections exert metabolic costs in *Drosophila melanogaster*. J. Exp. Biol. 228, jeb249424 (2025).

81. Liu, H., Wang, F., Cao, Y., Dang, Y. & Ge, B. The multifaceted functions of cGAS. J. Mol. Cell Biol. 14, mjac031 (2022).

82. Palmer, Clovis. S. Innate metabolic responses against viral infections. Nat. Metab. 4, 1245–1259 (2022).

83. Wang, J. & Meng, W. cGAS: Bridging Immunity and Metabolic Regulation. J. Mol. Cell Biol. mjaf018 (2025) doi:10.1093/jmcb/mjaf018.

84. West, A. P. et al. TLR signalling augments macrophage bactericidal activity through mitochondrial ROS. Nature 472, 476–480 (2011).

85. West, A. P., Shadel, G. S. & Ghosh, S. Mitochondria in innate immune responses. Nat. Rev. Immunol. 11, 389–402 (2011).

86. Marques, E., Kramer, R. & Ryan, D. G. Multifaceted mitochondria in innate immunity. Npj Metab. Health Dis. 2, 6 (2024).

87. Muta, T. & Iwanaga, S. The role of hemolymph coagulation in innate immunity. Curr. Opin. Immunol. 8, 41–47 (1996).

88. Ekengren, S. et al. A humoral stress response in Drosophila. Curr. Biol. 11, 714–718 (2001).

89. Vierstraete, E. et al. A proteomic approach for the analysis of instantly released wound and immune proteins in *Drosophila melanogaster* hemolymph. Proc. Natl. Acad. Sci. 101, 470–475 (2004).

90. Levy, F. et al. Peptidomic and proteomic analyses of the systemic immune response of Drosophila. Biochimie 86, 607–616 (2004).

91. Arrese, E. L. & Soulages, J. L. Insect Fat Body: Energy, Metabolism, and Regulation. Annu. Rev. Entomol. 55, 207–225 (2010).

92. Li, S., Yu, X. & Feng, Q. Fat Body Biology in the Last Decade. Annu. Rev. Entomol. 64, 315–333 (2019).

93. Krause, S. A., Overend, G., Dow, J. A. T. & Leader, D. P. FlyAtlas 2 in 2022: enhancements to the *Drosophila melanogaster* expression atlas. Nucleic Acids Res. 50, D1010–D1015 (2022).

94. Cohen, L. B., Lindsay, S. A., Xu, Y., Lin, S. J. H. & Wasserman, S. A. The Daisho Peptides Mediate Drosophila Defense Against a Subset of Filamentous Fungi. Front. Immunol. 11, 9 (2020).

95. Wiegand, I., Hilpert, K. & Hancock, R. E. W. Agar and broth dilution methods to determine the minimal inhibitory concentration (MIC) of antimicrobial substances. Nat. Protoc. 3, 163–175 (2008).

96. Kadeřábková, N., Mahmood, A. J. S. & Mavridou, D. A. I. Antibiotic susceptibility testing using minimum inhibitory concentration (MIC) assays. Npj Antimicrob. Resist. 2, 37 (2024).

97. Tattikota, S. G. et al. A single-cell survey of Drosophila blood. eLife 9, e54818 (2020).

98. Levashina, E. A. et al. Metchnikowin, a Novel Immune-Inducible Proline-Rich Peptide from *Drosophila* with Antibacterial and Antifungal Properties. Eur. J. Biochem. 233, 694–700 (1995).

99. Perlmutter, J. I., Chapman, J. R., Wilkinson, M. C., Nevarez-Saenz, I. & Unckless, R. L. A single amino acid polymorphism in natural Metchnikowin alleles of Drosophila results in systemic immunity and life history tradeoffs. PLOS Genet. 20, e1011155 (2024).

100. Patraquim, P., Magny, E. G., Pueyo, J. I., Platero, A. I. & Couso, J. P. Translation and natural selection of micropeptides from long non-canonical RNAs. Nat. Commun. 13, 6515 (2022).

101. Yang, H., Li, Q., Stroup, E. K., Wang, S. & Ji, Z. Widespread stable noncanonical peptides identified by integrated analyses of ribosome profiling and ORF features. Nat. Commun. 15, 1932 (2024).

102. Santos-Júnior, C. D., Pan, S., Zhao, X.-M. & Coelho, L. P. Macrel: antimicrobial peptide screening in genomes and metagenomes. PeerJ 8, e10555 (2020).

103. Mcclintock, B. Intranuclear systems controlling gene action and mutation. Brookhaven Symp. Biol. 58–74 (1956).

104. Britten, R. J. & Davidson, E. H. Repetitive and Non-Repetitive DNA Sequences and a Speculation on the Origins of Evolutionary Novelty. Q. Rev. Biol. 46, 111–138 (1971).

105. Faulkner, G. J. et al. The regulated retrotransposon transcriptome of mammalian cells. Nat. Genet. 41, 563–571 (2009).

106. Lisch, D. How important are transposons for plant evolution? Nat. Rev. Genet. 14, 49–61 (2013).

107. Bartonicek, N. et al. The retroelement Lx9 puts a brake on the immune response to virus infection. Nature 608, 757–765 (2022).

108. Lawlor, M. A., Cao, W. & Ellison, C. E. A transposon expression burst accompanies the activation of Y-chromosome fertility genes during Drosophila spermatogenesis. Nat. Commun. 12, 6854 (2021).

109. Fehlbaum, P. et al. Insect immunity. Septic injury of Drosophila induces the synthesis of a potent antifungal peptide with sequence homology to plant antifungal peptides. J. Biol. Chem. 269, 33159– 33163 (1994).

110. Landon, C., Sodano, P., Hetru, C., Hoffmann, J. & Ptak, M. Solution structure of drosomycin, the first inducible antifungal protein from insects. Protein Sci. 6, 1878–1884 (1997).

111. Fukuyama, H. et al. Landscape of protein-protein interactions in Drosophila immune deficiency signaling during bacterial challenge. Proc. Natl. Acad. Sci. U. S. A. 110, 10717–10722 (2013).

112. Duneau, D. F. et al. The Toll pathway underlies host sexual dimorphism in resistance to both Gram-negative and Gram-positive bacteria in mated Drosophila. BMC Biol. 15, 124 (2017).

113. Kaplan, E. L. & Meier, P. Nonparametric Estimation from Incomplete Observations. J. Am. Stat. Assoc. 53, 457–481 (1958).

114. Goel, M. K., Khanna, P. & Kishore, J. Understanding survival analysis: Kaplan-Meier estimate. Int. J. Ayurveda Res. 1, 274–278 (2010).

115. Abril-Pla, O. et al. PyMC: a modern, and comprehensive probabilistic programming framework in Python. PeerJ Comput. Sci. 9, e1516 (2023).

116. Kumar, R., Carroll, C., Hartikainen, A. & Martin, O. ArviZ a unified library for exploratory analysis of Bayesian models in Python. J. Open Source Softw. 4, 1143 (2019).

117. Kriegova, E. et al. PSMB2 and RPL32 are suitable denominators to normalize gene expression profiles in bronchoalveolar cells. BMC Mol. Biol. 9, 69 (2008).

118. Ponton, F., Chapuis, M.-P., Pernice, M., Sword, G. A. & Simpson, S. J. Evaluation of potential reference genes for reverse transcription-qPCR studies of physiological responses in Drosophila melanogaster. J. Insect Physiol. 57, 840–850 (2011).

119. FastQC(https://www.bioinformatics.babraham.ac.uk/projects/fastqc/).

120. Ewels, P., Magnusson, M., Lundin, S. & Käller, M. MultiQC: summarize analysis results for multiple tools and samples in a single report. Bioinformatics 32, 3047–3048 (2016).

121. Bolger, A. M., Lohse, M. & Usadel, B. Trimmomatic: a flexible trimmer for Illumina sequence data. Bioinformatics 30, 2114–2120 (2014).

122. Kim, D., Paggi, J. M., Park, C., Bennett, C. & Salzberg, S. L. Graph-based genome alignment and genotyping with HISAT2 and HISAT-genotype. Nat. Biotechnol. 37, 907–915 (2019).

123. Pertea, M. et al. StringTie enables improved reconstruction of a transcriptome from RNA-seq reads. Nat. Biotechnol. 33, 290–295 (2015).

124. Putri, G. H., Anders, S., Pyl, P. T., Pimanda, J. E. & Zanini, F. Analysing high-throughput sequencing data in Python with HTSeq 2.0. Bioinformatics 38, 2943–2945 (2022).

125. Love, M. I., Huber, W. & Anders, S. Moderated estimation of fold change and dispersion for RNA-seq data with DESeq2. Genome Biol. 15, 550 (2014).

126. Stephens, M. False discovery rates: a new deal. Biostatistics kxw041 (2016) doi:10.1093/biostatistics/kxw041.

127. Suvorov, A. et al. Widespread introgression across a phylogeny of 155 Drosophila genomes. Curr. Biol. 32, 111–123.e5 (2022).

128. Kumar, S. et al. TimeTree 5: An Expanded Resource for Species Divergence Times. Mol. Biol. Evol. 39, msac174 (2022).

129. Peng, J., Wang, B.-J., Svetec, N. & Zhao, L. Gene regulatory networks and essential transcription factors for de novo-originated genes. Nat. Ecol. Evol. 9, 1487–1498 (2025).

130. Jaccard, P. Étude comparative de la distribution florale dans une portion des Alpes et du Jura. 10.5169/SEALS-266450 (1901) doi:10.5169/SEALS-266450.

131. Mantel, N. The detection of disease clustering and a generalized regression approach. Cancer Res. 27, 209–220 (1967).

132. Li, H. et al. Fly Cell Atlas: A single-nucleus transcriptomic atlas of the adult fruit fly. Science 375, eabk2432 (2022).

133. ORFfinder(https://www.ncbi.nlm.nih.gov/orffinder/).

134. Teufel, F. et al. SignalP 6.0 predicts all five types of signal peptides using protein language models. Nat. Biotechnol. 40, 1023–1025 (2022).

135. Altschul, S. F., Gish, W., Miller, W., Myers, E. W. & Lipman, D. J. Basic local alignment search tool. J. Mol. Biol. 215, 403–410 (1990).

136. Birney, E., Clamp, M. & Durbin, R. GeneWise and Genomewise. Genome Res. 14, 988–995 (2004).

137. Katoh, K., Rozewicki, J. & Yamada, K. D. MAFFT online service: multiple sequence alignment, interactive sequence choice and visualization. Brief. Bioinform. 20, 1160–1166 (2019).

138. Suyama, M., Torrents, D. & Bork, P. PAL2NAL: robust conversion of protein sequence alignments into the corresponding codon alignments. Nucleic Acids Res. 34, W609–W612 (2006).

139. Nylander, J.A.A. MrModeltest Version 2. Program Distributed by the Author. (Evolutionary Biology Centre, Uppsala University, Uppsala.).

140. Huelsenbeck, J. P. & Ronquist, F. MRBAYES: Bayesian inference of phylogenetic trees. Bioinformatics 17, 754–755 (2001).

141. Gawde, U. et al. CAMPR4: a database of natural and synthetic antimicrobial peptides. Nucleic Acids Res. 51, D377–D383 (2023).

142. Veltri, D., Kamath, U. & Shehu, A. Deep learning improves antimicrobial peptide recognition. Bioinformatics 34, 2740–2747 (2018).

143. Wang, G., Li, X. & Wang, Z. APD3: the antimicrobial peptide database as a tool for research and education. Nucleic Acids Res. 44, D1087–D1093 (2016).

144. Ye, G. et al. LAMP2: a major update of the database linking antimicrobial peptides. Database 2020, baaa061 (2020).

145. The UniProt Consortium et al. UniProt: the Universal Protein Knowledgebase in 2023. Nucleic Acids Res. 51, D523–D531 (2023).

